# Receptor tyrosine kinase AXL regulates Golgi organization and function through an adhesion-Arf1 signaling axis in breast and lung cancers

**DOI:** 10.1101/2025.02.21.639453

**Authors:** Prachi Joshi, Arnav Saha, Radhika Malaviya, Debiprasad Panda, Grishma Mehta, Manojeet Pattanayak, Vibha Singh, Nagaraj Balasubramanian

## Abstract

Cell-matrix adhesion regulates membrane trafficking, Golgi organisation and function. Differential Golgi organisation in cancer cells could drive changes in trafficking and processing of cargoes. A simple screen evaluating Golgi organisation identifies breast (MDAMB231 vs MCF7) and lung cancer (A549 vs CaLu1) cell line pairs with differently organised Golgi, regulated differentially by loss of adhesion. In silico analysis of differentially expressed genes in the CCLE database, evaluated for their association with the Golgi in interaction networks and literature, identified AXL as a putative regulator of Golgi in these cancers. AXL prominently localized at the Golgi, undergoes displacement from the Golgi when inhibited by R428 and knocked down using siRNA, causing the Golgi to disorganize. AXL-dependent regulation of Golgi organisation is also dependent on cell-matrix adhesion. AXL binds active Arf1, whose recruitment to the Golgi is vital for its organisation. On loss of adhesion, loss of AXL and active Arf1 from the Golgi, drives its disorganization to affect Golgi-dependent microtubule acetylation and cell surface glycosylation in MDAMB231 and A549 cells respectively. Together, this validates our screen to identify novel regulators of the Golgi, and identifies AXL-Arf1 crosstalk as a vital mediator of its organisation and function in cancer cells.

## INTRODUCTION

Cell-matrix adhesion, essential for the survival, growth, and proliferation of all eukaryotic cells, is often deregulated by pathologies such as cancer (Berrier & Yamada, 2007; Reddig & Juliano, 2005). Cancer cells are unique in overcoming the requirement for cell-matrix adhesion, which promotes their anchorage-independent growth and metastasis(Janiszewska et al., 2020; Reddig & Juliano, 2005). Among the various alterations driving this anchorage independence in cancers are adhesion-independent growth signalling (Pawar et al., 2016) and changes in protein or cell surface glycosylation (Läubli & Borsig, 2019; Rambaruth & Dwek, 2011). Sustained activation or trafficking of growth receptors to the plasma membrane without adhesion promotes adhesion-independent signalling in cancers (Pawar et al., 2016; Schwartz, 1997). Alterations in glycosylation, on the other hand, support anchorage independence by overcoming the loss of adhesion-mediated cell death (anoikis) and by evasion of death ligands (Petrosyan et al., 2014; Piyush et al., 2017). Glycan alterations are mainly driven by changes in the expression or localisation of glycosylation-associated proteins and Golgi organisation, which is the central hub for glycosylation reactions (Bhat et al., 2017; Petrosyan, 2015).

The Golgi apparatus comprises multiple membranous sacs called cisternae, which stack up in a defined sequence (cis-, medial and trans-Golgi). In mammalian cells, lateral linking between the stacks forms a Golgi ribbon localised to the perinuclear region (Marsh & Howell, 2002; Nakamura et al., 2012; Rambourg & Clermont, 1997). Together with the ER-Golgi intermediate compartment (ERGIC) and the trans-Golgi network (TGN), the Golgi functions as a complex and dynamic system, which is the site for post-translational modifications (PTMs) and trafficking events (Huang & Wang, 2017; Ward et al., 2001). These Golgi functions depend on several Golgi-associated regulatory pathways, including the intact organisation of the Golgi ribbon (Petrosyan, 2019; Stanley, 2011; X. Zhang & Wang, 2015).

Alterations to these regulatory pathways or the Golgi organisation affect the trafficking and glycosylation reactions at the Golgi, which could, in turn, facilitate cancer progression (Petrosyan, 2015). Golgi organisation is often reported to be inherently disorganised in several types of cancers (Kellokumpu et al., 2002; Petrosyan et al., 2014; Rivinoja et al., 2006). However, the role and regulation of altered Golgi organisation and its function in cancers is only recently being explored and remains to be completely understood (Bajaj et al., 2022; Bhat et al., 2017; Bui et al., 2021; Howley & Howe, 2018).

While Golgi organisation and adhesion-dependent signalling are known to be altered in cancer cells, earlier studies from our lab have shown that adhesion can regulate Golgi organisation (Singh et al., 2018). In non-transformed cells, loss of adhesion-mediated Golgi disorganisation is mediated by the inactivation of the small GTPase Arf1, leading to Dynein motor protein disengagement from the Golgi membranes. Distinct changes in cell surface glycosylation accompanied the observed Golgi disorganisation in these non-adherent cells. This suggests the presence of cell adhesion – Arf1 – Dynein – Golgi Organization – Golgi function pathway in normal cells (Singh et al., 2018). If and how this adhesion-mediated regulation of Golgi is perturbed in cancers, which are inherently anchorage-independent, remains unknown. Differential regulation of the Golgi in cancers could further impact the Golgi function and support cancer progression. Golgi organisation in cancers is variable, with some cancer cell lines seen to have ’dispersed’ or ’fragmented’ Golgi, while others have ’normal’ intact Golgi (Bhat et al., 2017; Petrosyan, 2015; Petrosyan et al., 2014). Amongst the known regulators of Golgi organisation, which could potentially be perturbed in cancers, are Golgi matrix proteins (Witkos & Lowe, 2015; Xiang & Wang, 2011; X. Zhang & Wang, 2015), Golgi-associated GTPases (Goud et al., 2018; Thomas & Fromme, 2020; Ward et al., 2001), cytoskeletal proteins (Egea et al., 2015; Kulkarni-Gosavi et al., 2019), and kinases(Chia et al., 2012; Kimura et al., 2018; Mao et al., 2013; Weller et al., 2010). Cancer cells of similar origin with different Golgi organisations could constitute a unique system to evaluate this regulation and possibly identify novel regulators that could mediate it. Our simple screen in breast (MDAMB231 and MCF7) and lung cancer (A549 and Calu1) cell lines with organised vs dispersed Golgi allows us to evaluate differentially expressed genes as possible candidate regulators of the Golgi.

This led us to AXL, a high-score candidate we shortlisted in both breast and lung cancer cell lines. AXL is a transmembrane receptor tyrosine kinase from the TAM family of receptors, which plays a significant role in multiple cellular processes, including cell growth, proliferation, cell survival, cell adhesion, and apoptosis (Auyez et al., 2021; Wium et al., 2021). AXL was first discovered as an oncogene, and since then, its role in cancer progression has been extensively researched. Overexpression of AXL has been shown in several human malignancies, including breast cancer (Holland et al., 2010; Zajac et al., 2020), acute myeloid leukaemia (Hong et al., 2008), non-small cell lung cancer (NSCLC) (Iida et al., 2017; Z. Zhang et al., 2012), ovarian cancer (Kanlikilicer et al., 2017), glioblastoma (Onken et al., 2016), and neuroblastoma (Debruyne et al., 2016). Altered expression or activation of AXL in these cancers promotes invasiveness and metastasis, epithelial-to-mesenchymal transition, drug resistance, and anchorage-independent growth (Debruyne et al., 2016; Lay et al., 2007; Scaltriti et al., 2016; Taniguchi et al., 2019; Ye et al., 2010). AXL is activated by its ligand Gas6, which belongs to the vitamin K-dependent family of proteins. Gas6 and AXL bind at a one-to-one receptor-to-ligand ratio and then dimerise with another Gas6–AXL complex, promoting trans-autophosphorylation to initiate downstream signalling (Tanaka & Siemann, 2021). Amongst the six phosphorylation sites on AXL, the three C-terminal sites - Tyr698, Tyr702, and Tyr703 are relatively conserved among the TAM receptors and are indispensable for the complete functions of the kinase (Auyez et al., 2021; Zhu et al., 2019).

Few studies have explored a possible crosstalk between AXL and the Golgi apparatus. A recent study showed that inhibition of AXL affects its localisation at the Golgi and impairs directed migration in breast cancer cells (Hs578t)(Zajac et al., 2020). A screen for regulators of the Golgi organisation targeting the kinome and phosphotome in HeLa cells also identified AXL as one of the possible regulators of the Golgi organisation. Knockdown of AXL in this study was shown to cause the Golgi to become more condensed in HeLa cells (Chia et al., 2012). In their study, Haines et al. suggested that AXL could bind the Golgi-associated GTPase Arf1 in breast cancer cells (Haines et al., 2015). These observations point to AXL as a potential regulator at the Golgi, which could be deregulated in cancers to affect Golgi organisation and function.

Investigating differences in Golgi organisation in breast and lung cancer cells, our study identifies AXL as a vital regulator of Golgi organisation and function in cancer cells. Its role in regulating the loss of adhesion-mediated Golgi organisation also implicates AXL in a more physiological setting. The differential recruitment of AXL at the Golgi and crosstalk with Arf1 could hence play a vital role in Golgi organisation and function.

## MATERIALS AND METHODS

### Reagents

Accutase (Cat. No. #A6964), Fibronectin (Cat. No. #F2006) and Triton-X 100 (Cat. No. #T8787) were purchased from Sigma. Lipofectamine 2000 was purchased from Invitrogen (Cat. No. # 11668019). Fluorophore conjugated lectin probes were purchased from Invitrogen Molecular probes - ConA-Alexa488 (Cat. No. #C11252), WGA-Alexa488 (Cat. No. #W11261), PNA-Alexa488 (Cat. No. #L21409). DMSO was purchased from Sigma (Cat. No. # D2438). Bemcentinib (R428) was purchased from Cayman USA (Cat. No. #21523) and Golgicide A (Cat. No. #G0923) was purchased from Sigma. Immobilon Western Chemiluminescence substrate was purchased from Millipore (Cat. No. #WBKLS0500). Trizol was purchased from Ambion (Cat. No. # 15596018). Fluoramount-G was purchased from Southern Biotech (Cat. No. #0100-01). cDNA synthesis kit (Takara Cat. No. #6210A) was obtained from Dr. Nishad Matange, IISER Pune. SYBR green mix was purchased from Takara (Cat. No. #RR820A). BCA protein estimation kit (Cat. No. #23225) was purchased from Pierce. DAPI was purchased from Merck (Cat. No. 5.08741.0001)

### Antibodies

pAkt S473 (1:1000) (Cat. No. #4060S), Akt (1:2000) (Cat. No. # 9272S). AXL Clone C89E7 (1:2000 for western blotting and 1:400 for immunofluorescence assay) (Cat. No. #8661S) and pAXL D12B2 (Y702) (1:500) (Cat. No. #5724S) was purchased from Cell Signaling. GAPDH (1:5000) (Cat. No. #9545) was purchased from Sigma. GM130 Clone 35 (1:250 for western blotting and 1:100 for immunofluorescence assay), (Cat. No. # 612008) was purchased from BD Transduction. Arf1 Clone 1D9 (1:500), (Cat. No. # MA3-060) was purchased from Invitrogen. β-tubulin Clone E7 (1:2000), (Cat. No. #AB_2315513) and Alpha Tubulin Clone 12G10 (1:200), (Cat no. #AB_1157911) were purchased from Developmental Studies Hybridoma Bank (DSHB). Anti-Acetylated Tubulin (1:2000), (Cat. No. T7451) was purchased from Sigma. Anti-GBF1 antibody (1:1000) (Cat. No. ab86071) was purchased from Abcam. Secondary Fluorescent conjugated antibodies (Alexa488 and Alexa568) were purchased from Invitrogen Molecular Probes and used at a dilution of 1:1000. HRP-conjugated secondary antibodies were purchased from Jackson Immunoresearch and used at a dilution of 1:5000.

### Plasmids and Oligos

GalTase-RFP and Mannosidase-GFP constructs were obtained from Dr. Jennifer Lippincott Schwartz (HHMI). ABD-GFP and ABD-RFP constructs were obtained from Dr. Satyajit Mayor, National Centre for Biological Sciences, Bangalore (NCBS), India. mCherry-tagged Arf1-WT and Arf1-Q71L constructs were made by releasing the Arf1 gene from GFP constructs and cloning the same into an empty mCherry-N1 vector. Primers used for RT-PCR experiments were designed using a Primer design tool from Integrated DNA Technologies and were ordered from Eurofins. Primer sequence for AXL –

Forward – GTCTAGCTGACCGTGTCTAC

Reverse – CCTGGCGCAGATAGTCATAA.

AXL siRNA sequences confirmed in earlier studies (Holland et al., 2010) were purchased from Sigma oligos.

### siAXL#1

Forward – GAAAGAAGGAGACCCGTTA

Reverse – TAACGGGTCTCCTTCTTTC

### siAXL#2

Forward – CCAAGAAGATCTACAATGG

Reverse – CCATTGTAGATCTTCTTGG

All sequences were checked and confirmed for target specificity using the NCBI nucleotide BLAST tool.

### Cell culture and transfections

MDAMB231 and CaLu1 cell lines were obtained from ECACC. A549 cell line was obtained from ATCC. The MCF7 cell line was obtained from Prof. Sanjay Gupta at ACTREC, Navi Mumbai, India. All the above cell lines were cultured using Gibco DMEM from Thermo Fisher Scientific, with the addition of 10% Penstrep and 5% FBS. 0.05% Trypsin was used to detach cells, and an excess culturing medium was used to neutralise the action of trypsin. BEAS2B cell line was obtained from Prof. Shantanu Chaudhary’s lab at IGIB, New Delhi and the cells were cultured using LHC9 media. 0.05% Trypsin was used to detach cells and Trypsin inhibitor purchased from Roche (Cat. No. #10109886001) was used to neutralize trypsin. MCF10A cell line was obtained from Dr. Madhura Kulkarni’s lab at CTCR, IISER Pune & Prashanti Cancer Care Mission (PCCM), and the cells were cultured using Gibco DMEM/F12 media (Cat. No. 12500062) with the addition of 5% horse serum (Invitrogen#16050-122), 10% Penstrep, 20 ng/ml EGF, 0.5 mg/ml hydrocortisone (Sigma Cat. No. H0888-5G), 100 ng/ml cholera toxin and 10 ug/ml (Sigma Cat. No. C8052-1MG) and Insulin (Sigma Cat. No. I1882-100MG). For transfection studies, cells were seeded in 6 cm dishes to attain a confluency of 60% and allowed to attach and spread for 5 hours. Using Gibco OptiMEM medium and transfection agent Lipofectamine 2000 from Thermo Fisher Scientific, the transfection mix was prepared and kept at room temperature for 30 minutes before adding to the cells seeded in 6cm dishes. Media in transfected dishes was changed 12 hours post-transfection, and cells were used for experiments 36 hours post-transfection.

### Suspension assay experiments

Cell lines maintained in their respective media were grown up to 75% confluency in 10cm or 6cm dishes, as required. 0.05% Trypsin or Accutase (for lectin labelling experiments) was washed out using excess media to detach cells. One aliquot was kept aside from collected cells and processed for the 5-minute (just detached) timepoint if required. For the suspension assay, cells were gently mixed with a culture medium containing 1% methylcellulose and incubated at 37°C with 5% CO_2_ for a specified duration. Post-suspension, methyl cellulose was washed out using culture medium, and 4°C conditions were maintained throughout to harvest cells from suspension. Collected cells were distributed to suspension and re-adhesion time points as required. For re-adhesion time points, cells post-suspension was seeded on 22 x 22mm coverslips or 6cm dishes pre-coated with 2 mg/ml or 10 mg/ml Fibronectin, respectively. For inhibition experiments, respective inhibitors were added to the suspension mix, media washes and re-adhesion timepoints. Each timepoint was then processed for lectin labelling, fixed with PFA for IFA/imaging, or kept at -80°C for preparing cell lysates and for active Arf1 pulldown experiments. Cell numbers used for suspension assays were variable depending on the experiment.

### Lectin labelling and Flow Cytometry analysis

Cells harvested from suspension were given a PBS wash to remove residual media, then added to a lectin labelling mix containing fluorophore-conjugated lectin diluted in 1X PBS at optimized lectin concentration. The labelling reaction was kept in ice under dark conditions for 15 minutes, followed by two washes with cold 1X PBS. PFA (3.5%) was used to fix the lectin-labelled cell samples, which were then resuspended in cold 1X PBS (350 ul) for Flow cytometry analysis. Samples were run on BD Celesta Flow Cytometer and data analysis was done using BD FACS Diva software. A morphologically uniform cell population presented by the forward and side scatter (FSC and SSC) plot was selected using polygon gating for data acquisition. A maximum of 15000 events were recorded in the gated area for every sample, and the median fluorescent intensity obtained for the selected population was used.

### Immunofluorescence assay (IFA)

For IFA, cells fixed with PFA were incubated in a Permeabilization buffer for 15 minutes at room temperature. A permeabilization buffer was made with Triton X 100 (0.05%) diluted in 5% BSA and made in 1X PBS. Post permeabilization, two washes were given with 1XPBS followed by blocking with 5% BSA at room temperature for 30 min to 1 hr. Two washes were given after blocking. Samples were incubated with Primary antibody overnight at 4°C. Three washes were given post-incubation with primary antibody followed by a 1-hour incubation with secondary fluorescent antibody (1:1000 dilution in 5% BSA) at room temperature. Samples were given three washes and then mounted on slides using Fluoramount-G. Slides were maintained at room temperature under dark conditions to dry appropriately, then moved to 4°C until confocal imaging.

### Inhibitor studies

Cells grown to 75% confluency were treated with increasing concentrations of AXL inhibitor R428 (0.5 μM, 1 μM, 2 μM) for 12 hours before using cells for preparing lysates or preparing slides for confocal imaging. Optimal 1 μM R428 concentration was chosen similarly used to treat cells for 12 hrs before being held in suspension with the inhibitor and further processed for lysis or confocal imaging. Cells were incubated with 1 μM of R428 for increasing timepoints (10 min, 20 min, 30 min, 40 min, 60 min and 12 hrs) to evaluate its effect in time kinetic studies. For experiments inhibiting Arf GEFs, cells were suspended for 90 minutes and then treated with 0.5 or 1μM of GCA for 30 min in suspension. Adherent cells were treated with GCA (0.5 μM/1 μM) for 30 minutes and lysed or fixed as required. In experiments using R428 or GCA, DMSO was used as a control. During these experiments, respective inhibitors were included in all incubation steps of the protocol. In suspension assays inhibitors were also included in the washes between incubation and lysis or fixing.

### siRNA mediated knockdowns

For siRNA mediated knockdowns, cells were seeded in 3.5 cm dishes to attain 50% confluency and allowed to attach and spread for 4 hrs. Transfection mix was prepared using Gibco OptiMEM, transfection reagent RNAi MAX and 50 pmol of siAXL and allowed to incubate at room temperature for 30 min before adding to the cells seeded on 3.5 cm dishes (5 x 10^5^ cells for MDAMB231 and 4 x 10^5^ cells for A549). Second shot of siRNA was given to cells after 24 hrs. Cells were used for experiments after 48 hrs post-second shot. Control cells were treated with only RNAi MAX.

### Arf1 activity assay

Activity assay was performed simultaneously for all time points/samples in an experiment to ensure that the same amount of beads was used to compare the samples. Samples for pulldown assay were processed immediately at respective time points or frozen at -80°C before performing activity assay.

Active Arf1 pulldown was performed using Glutathione Sepharose A beads bound to 60 ug of GST-GGA3. Samples were lysed with 500 ul activity assay buffer of which 400 ul were incubated with beads at 4°C for 35 min at 9 rpm. The remaining 100 ul of lysate was mixed with 5X Laemmli buffer and used as the whole cell lysate fraction. Post incubation with beads samples were washed and GST-GGA3 bound Arf1 eluted with 1X Laemmli buffer. Samples were boiled at 95°C and given a short spin before performing Western blotting. Blots were incubated with required antibodies developed using Immobilon reagents and analysed using ImageJ software (described under the section ‘SDS-PAGE and Western blotting).

To determine the percentage activity of Arf1, the following calculation was used as described in earlier studies.

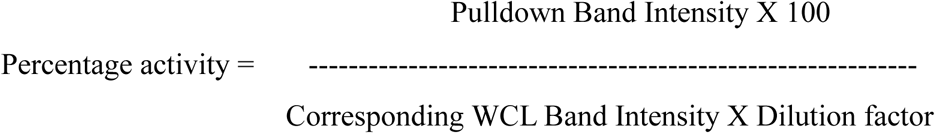

### Protein estimation with BCA kit

Samples were lysed in RIPA buffer with freshly added protease inhibitors and kept in ice for 30 minutes of lysis. Lysates were then spun down at 14000 rpm; 4°C for 15min. Supernatant was collected in a fresh tube as the lysate. 10ul of samples were set aside for BCA, and the rest was lysed in Laemmli buffer and boiled at 95°C, followed by a short spin. Lysates were stored at -20°C until gel run and western. Protein estimation was done as follows: in a 96-well plate, unknown samples diluted to 1:5 times with RIPA buffer were loaded in triplicates along with freshly prepared standards for BSA diluted in RIPA buffer (range – 0 mg/ml to 2 mg/ml). Working reagent from the Pierce BCA kit was prepared by mixing Reagent A and Reagent B in a ratio of 50:1. 200ul of working reagent was added to each well containing 10ul of sample or BSA standard. The plate was incubated at 37°C for 30 minutes and then scanned at 562 nm to determine absorbance using a plate reader (PerkinElmer, Ensight). Absorbance values for BSA concentrations were plotted to obtain a standard curve, which determined the concentration for the unknown sample.

### SDS-PAGE and Western Blotting

Required amount of protein (20 ug, 30 ug) or cell equivalent amount of lysate was loaded in gels for SDS-PAGE. Protein samples run on gels were transferred to the PVDF membrane using a methanol-containing buffer or sodium bicarbonate buffer. Post transfer, the PVDF membrane was blocked in 5% skimmed milk (made in 1X TBS-Tween 20 (0.1%)) at room temperature for 1 hr at 10 rpm. The membrane was washed with 1X TBST; then, individual blots were incubated in respective Primary antibody dilutions in 5% BSA in 1X TBST overnight at 4°C. Blots were washed three times with 1X TBST, followed by incubation at RT with respective HRP-conjugated secondary antibody solutions made in 2.5% skimmed milk in 1X TBST. After this incubation blots were washed thrice with 1X TBST and developed with direct or diluted (with 1X TBST) Immobilon substrate. Blot images were captured using LAS4000 Chemiluminescent Imager. Densitometric analysis was done using ImageJ FIJI software.

### Anchorage-Independent growth (AIG) assay

Cells grown to 75% confluency were treated with DMSO or R428 (1μM) for 12 hourse before performing anchorage-independent growth assay. Detached cells were counted and 5000 cells from control and R428 treated cells each, were mixed with 0.3% agar containing DMEM and layered on top of 0.5% agar base per well in 6-well plates. Each of the control and inhibitor-treated cells were plated in duplicates. The agar was allowed to solidify, and 1.5 mL of culture medium with DMSO or R428 was added. These dishes were maintained in the incubator for 15 days and DMSO or R428 was added freshly when culture medium was changed every three days. The colonies formed in agar at the end of the 15-day incubation were stained with 0.05% crystal violet dissolved in 20% ethanol for 1 hour at room temperature and de-stained by repeated washing with distilled water until stained colonies were visible. The colonies were then imaged on an Olympus MVXC10 microscope at 1X zoom with 1X objective in the HDR mode and counted using the particle analysis tool of Image J software.

### Confocal imaging

Slides were imaged using the ZEISS confocal (LSM710) using a 63X oil immersion objective. For profiling Golgi organisation, cells observed at confocal were categorised manually as being with organised Golgi or disorganised Golgi. For ABD-RFP profiling studies, cells showing localisation of ABD-RFP at the Golgi vs cells without were identified and counted. A minimum of 100 and a maximum of 200 cells were counted for each experimental treatment and categorised according to their phenotype to arrive at a percent population of cells in either category. Images were captured with an average of 4, zoomed at 5X / 6X for suspended cells and 2X for stable adherent cells. A scan speed of 5 for cross-sectional images and 7 for obtaining z-stacks was used. Leica DM6 was used for evaluating active Arf1 localisation using an ABD-RFP marker expression in cells.

### Image Analysis and quantitation

ImageJ FIJI was used for all imaging experiments to process images with scale bars until otherwise specified. Overlap of markers in images when looked at using a line plot was done using the ImageJ FIJI software. Huygens Professional software (SVI) was used to deconvolution LSM files. Deconvoluted images were used to generate Maximum Intensity Projection (MIP) images and 3D Surface Rendered (SR) images of the Golgi using the Visualisation tool in Huygens. Colocalization analysis for data showing AXL localisation at the Golgi was performed using the Huygens software ’Colocalization Analyzer Advanced’ tool. Costes’ method was used to set a threshold for each image, and the GM130 (Alexa488) channel was manually thresholded and analysed by the software to obtain Pearson’s coefficient value.

### Gene expression analysis for regulators of Golgi organisation (In-silico study)

A comprehensive listing of genes associated with the Golgi apparatus was done using the NCBI Gene Ontology (GO) tool. A search using the terms’ Golgi organisation human’ yielded 131 human genes associated with the term Golgi organisation. Gene expression data (RPKM) was obtained from the CCLE database using the UCSC Xena browser for MDAMB231, MCF7, A549 and CaLu1 cell lines. Fold change in gene expression was calculated between the pair of breast cancer (BC) cell lines and the pair of lung cancer (LC) cell lines independently, and a difference of ≥10 (BC) ≥5 (LC) was considered significant to obtain the differentially expressed genes (DEGs). This cut off was set to ensure that the number of genes picked is in the top 50. STRING-DB version 11.0 created a primary protein interaction network for each DEG. No restriction was applied over the number of interactors displayed in the network to cover all possible primary interactions. Interactions based on experimental evidence with a confidence score of ≥0.4 were considered in the study. Each of the primary interactors obtained was evaluated for any known role in the regulation of Golgi organisation or Golgi function.

Scoring of genes - Knowing the Golgi organisation phenotype in the selected cell lines, the primary list of selected genes was scored based on available data in the literature for the effect of gene knockdown on Golgi organisation. Accordingly, genes whose knockdown affected the Golgi phenotype comparable to a drop in levels observed in cancer cell lines were scored as 2. Those that are involved with the Golgi in literature studies but whose expression did not agree with their knockdown Golgi phenotype from literature were scored as 1. Genes having no available data to suggest their knockdown affects Golgi organisation were scored as 0. A second score was given based on the number of primary interactors obtained from STRING analysis that have a direct role in Golgi’s organisation or function. The two scores were added for each gene, and the DEGs with a combined score of 3 or higher were selected for further evaluation. This gave us a shortlist of 20 DEGs for breast cancer and 14 DEGs for lung cancer. Amongst them 8 genes were common between both breast and lung cancer. Having looked at genes with the top 3 scores in this list of 8, we chose AXL to evaluate its role in regulating Golgi organisation and function in these cancers.

### RNA isolation and cDNA synthesis

For preparing RNA samples, 8×10^5^ cells were seeded for 14 hrs were lysed in Trizol (Ambion), followed by RNA isolation using the PCI method (Phenol, chloroform, and Isopropyl alcohol). After ethanol washes, the RNA was allowed to dry for 15 minutes at 37°C and resuspended in 50 ul of Nuclease Free Water (NFW). RNA concentration and purity were checked using NanoDrop 2000. 1 ug of RNA was mixed with Gel loading buffer and run on 10% Agarose gels to check the quality of RNA samples. These RNA sample stocks were stored at -80°C. Using the PrimeScript cDNA synthesis kit (Takara), cDNA samples were prepared from at least five sets of RNA (1ug) for each MDAMB231, MCF7, A549 and CaLu1 cell lines. Details of the PCR cycle run were as follows - 25°C for 2min, 25°C for 15 seconds, 37°C for 25 min, 85°C for 1min 30seconds. cDNA samples were stored at -20°C for Real-time PCR studies.

### Real-Time PCR

Primers were tested for efficiency by running RT-PCR for a range of cDNA dilutions. Primer efficiency was determined by three criteria – single peak for melt curve, Ct value for the highest concentration of cDNA, and slope value for a range of cDNA concentrations. Post RT-PCR, the product was run on Agarose gels to confirm if the product ran at the expected band size. All RT-PCR experiments for comparative gene expression analysis were performed using 1:1 dilution of cDNA. SYBR green mix (Takara) was used to detect and quantify gene expression. Actin was used as the standard control. 5 ul reaction mix was loaded in triplicates for each sample. The plate was given a quick spin before running the RT-PCR cycle, the protocol for which was as follows: 95°C for 3 minutes (once per sample). Plate reading was recorded after the two steps of 95°C, 10 seconds – 60°C; 25 seconds; this cycle was repeated 40 times per sample. Recorded data was analysed for Delta Ct and Fold change. The relative gene expression between MDAMB231 and MCF7 or A549 and CaLu1 cell lines was compared to confirm the differential expression of AXL.

### Statistics

All the statistical analysis was done using GraphPad Prism analysis software. Statistical analysis for western blotting using absolute data (not normalised to a condition or control) was done using the two-tailed unpaired Mann-Whitney U test. For western blot comparisons data normalised to control or a certain experimental condition, the two-tailed single sample t-test was used. Statistical analysis for Golgi profiling data, comparing the percentage of cells with a specific Golgi phenotype at different experimental or treatment conditions, was done using one-way ANOVA, a multiple comparison test, with Tukey’s method for error correction. The Pearson’s coefficient analysis data for localisation of AXL at the Golgi was tested for statistical significance using one-way ANOVA, a multiple comparison test with Tukey’s method for error correction.

## RESULTS

### Adhesion independent regulation of Golgi organisation in cancers

Cell-matrix adhesion mediated signalling regulates Golgi organisation in anchorage-dependent cells (Singh et al., 2018). Cancer cells in being anchorage-independent do not require adhesion-dependent signalling for growth and survival. This is seen in the de-regulation of many adhesion-dependent pathways, prompting us to ask, what about the Golgi? A simple screen for Golgi organisation across multiple cancer cell lines using a trans-Golgi marker (GalTase-RFP) observed much variation across cell lines. Breast cancer MDAMB231 and lung cancer A549 cells showed a predominantly organised Golgi phenotype, while in breast cancer MCF7 and lung cancer CaLu1 cells the Golgi is predominantly disorganised **(Fig 1A)**. On loss of adhesion MCF7 **(Fig 1C)** and CaLu1 **(Fig 1E)** cells retained their disorganised Golgi phenotype, detected using the cis-medial (MannosidaseII-GFP) and trans-Golgi marker (GalTase-RFP). In A549 cells, the Golgi stays intact on loss of adhesion **(Fig 1D)**, unlike the non-transformed lung epithelial BEAS2B cells where it is dispersed **(Supp Figure 1A)**. In MDAMB231 cells the Golgi has a ’normal’ dispersed phenotype **(Fig 1B)** as seen in the non-transformed breast epithelial MCF10A cells on loss of adhesion **(Supp Figure 1B, 1C)**. This indicates that adhesion-dependent regulation of Golgi is also highly variable in different cancer cells, even between cell lines of similar tissue origins. We hence asked how these differences in Golgi organisations could affect cancer cell function and what regulatory pathway(s) downstream of cell-matrix adhesion could mediate this.

**Figure 1:**
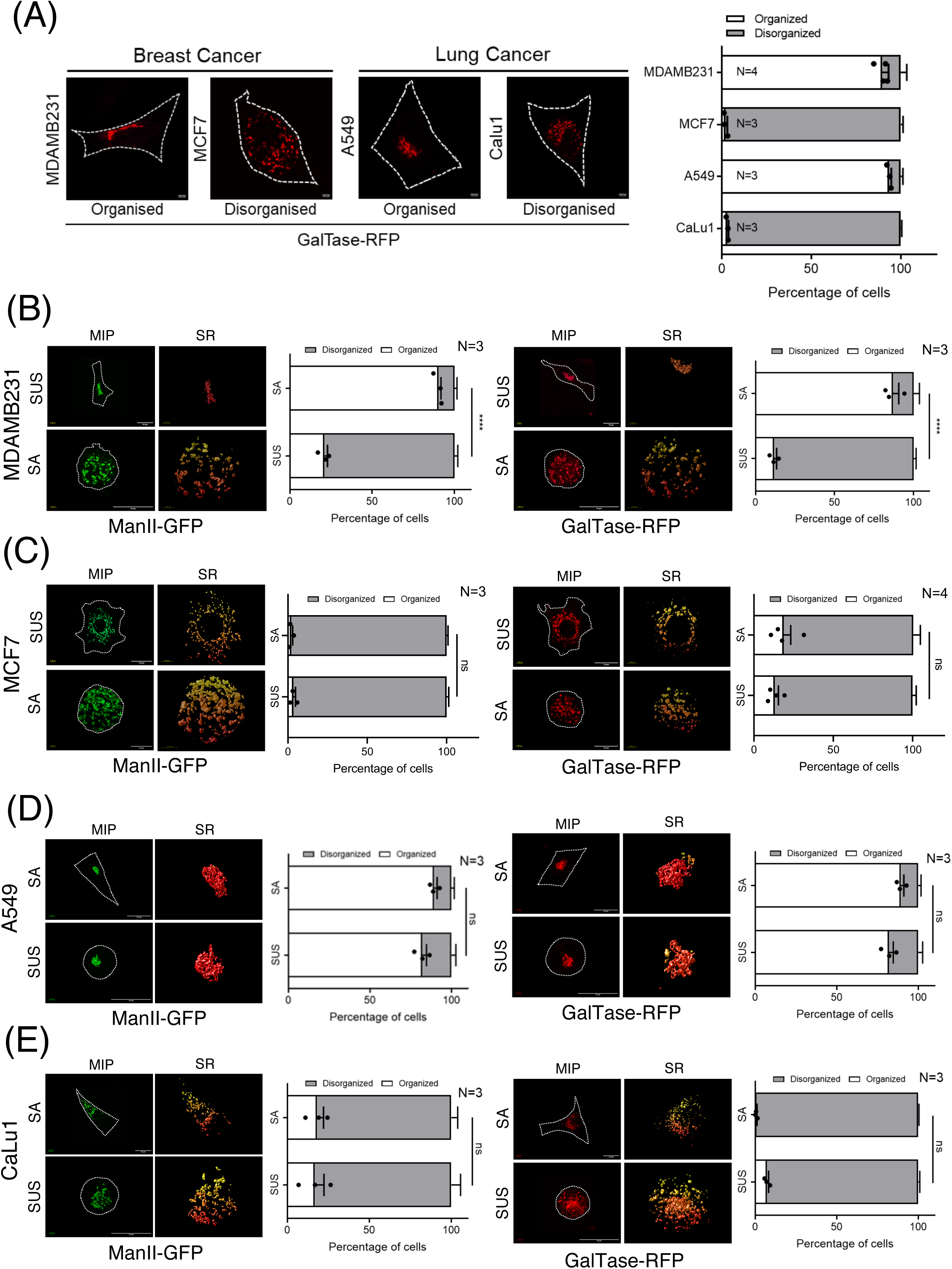
Adhesion independent regulation of Golgi organization in breast and lung cancer cells. **(A)** Stable adherent MDAMB231, MCF7, A549 and CaLu1 cells transfected with GalTase-RFP (trans-Golgi marker). Representative cross-sectional images show the predominant Golgi organization phenotype. Percentage distribution profile of cells (n ≥200) show organized (white) and disorganized (grey) Golgi in cells. Graph represents mean±SEM from three-four independent experiments. **(B-E)** Stable adherent (SA) and non-adherent (SUS) MDAMB231, MCF7, A549 and CaLu1 cells transfected with ManII-GFP (cis-medial Golgi marker) or GalTase-RFP (trans-Golgi marker). Representative deconvoluted Z-stacks images show the predominant phenotype as maximum intensity projection (MIP) and a zoomed image of the Golgi with surface rendering (SR). Percentage distribution profiles of cells (n ≥200) show organized (white) and disorganized (grey) Golgi in cells. Graphs represent mean±SEM of percentage distribution from three-four independent experiments. Statistical analysis done using one-way ANOVA multiple comparisons test with Tukey’s method for error correction. Scale bars shown are 4.22 µm in **(A)** and 10 µm in **(B-E)**. (*p≤0.05, **p ≤ 0.01, ***p ≤ 0.001, ****p ≤ 0.0001, ns= not significant).

### Differentially expressed genes in breast and lung cancer cells as potential regulators of Golgi organisation

Differences in Golgi organisation seen in breast and lung cancer cells, regulated by adhesion could be mediated by differences in gene expression of Golgi-associated or Golgi-regulatory proteins. To evaluate this, we designed an *in-silico* comparative evaluation of differentially expressed genes in MDAMB231 vs MCF7, and A549 vs CaLu1 **(Fig 2A)**. Using the NCBI Gene Ontology database and a collated list of known regulators of Golgi organisation and/or function reported in literature we arrived at a list of 390 possible genes that could affect Golgi organisation. CCLE database was used to compare the differential mRNA expression of these regulators between MDAMB231 vs MCF7 and A549 vs CaLu1 cells. This revealed 42 differentially expressed genes (DEGs) in breast cancer and 35 DEGs in lung cancer that show a >10 and > 5-fold change in expression respectively. The fold change cut-off applied, was adjusted for breast and lung cancer such that the genes to be evaluated were in the top 50. For further evaluation, a scoring system was applied to these shortlisted DEGs. The first component of the score is based on the effect knockdown of the gene has on Golgi organisation in existing literature. The second component of this score was based on the number of primary interactors of the gene of interest in the STRING database that are involved in regulating Golgi organisation or function. Setting a cut-off of 3 or more for this combined score, we ranked the shortlisted 20 genes (breast cancer) and 15 genes (lung cancer) as candidate regulators for testing their role in differential Golgi organisation **(Fig 2A)**. Of these, 8 shortlisted genes were common between breast and lung cancer cells. Among the top 3 candidates - AXL was chosen for this study while we are now evaluating the other genes **(Fig. 2B)**.

**Figure 2.**
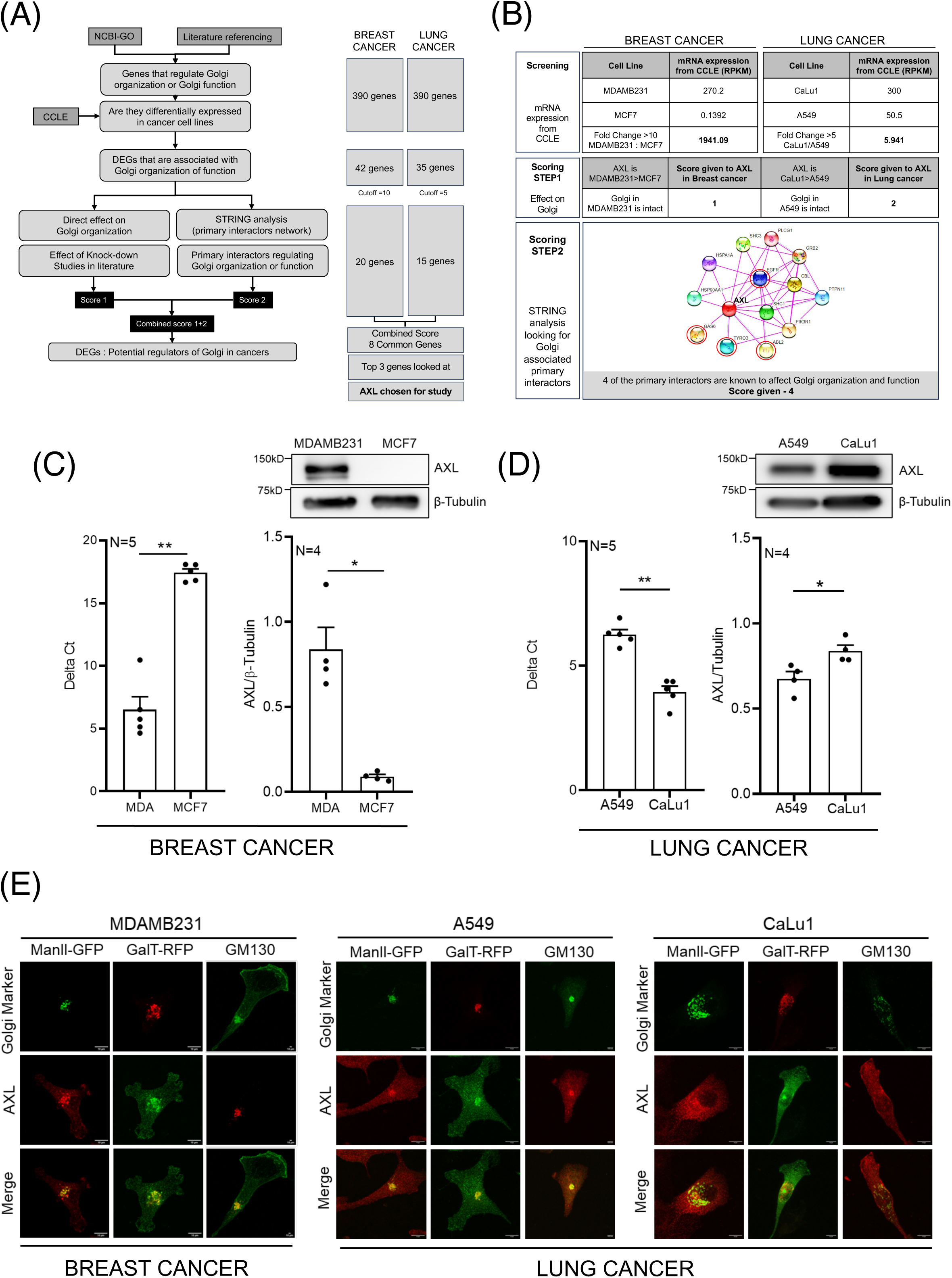
Differentially expressed genes in breast and lung cancer cells. **(A)** Schematic representation shows the sequence of steps followed in the *in-silico* analysis. The number of shortlisted genes at each stage of the protocol for breast and lung cancer cells are also shown. **(B)** Details of the *in-silico* analysis for AXL in breast and lung cancer cell line pairs showing their screening based on mRNA expression in CCLE database (Screening), scores given based on their known effect on Golgi organization (Scoring STEP1) and based on number of primary interactors known to be associated with the Golgi derived using STRING analysis (Scoring STEP2). **(C, D)** Comparative mRNA and protein expression of AXL in **(C)** MDAMB231 and MCF7 (BREAST CANCER) and **(D)** A549 vs CaLu1 cells (LUNG CANCER). Graph represents mean±SEM of delta Ct values obtained from qPCR analysis from five independent experiments (shown on left) and ratio of western blot densitometric band intensities for AXL and β-Tubulin shown as mean±SEM from four independent experiments. Representative western blots are shown above the graph. **(E)** Representative cross-sectional images for AXL localization at the Golgi with Golgi markers, GM130 (red in MDAMB231 and green in A549 and CaLu1 cells) or ManII-GFP (green) or GalTase-RFP (red) in MDAMB231 (BREAST CANCER) or A549 and CaLu1 cells (LUNG CANCER). Top panel shows the Golgi marker, followed by AXL and the last panel shows merged images. Statistical analysis was done using Mann-Whitney U test. All scale bars shown are 10 µm. (*p≤0.05, **p ≤ 0.01, ***p ≤ 0.001, ****p ≤ 0.0001, ns= not significant).

We tested AXL’s mRNA and protein expression in these cell lines **(Fig 2C**, **Fig 2D)**. In concurrence with the CCLE data, AXL expression was higher in MDAMB231 cells than MCF7 cells and in CaLu1 cells than in A549 cells. MCF7 cells have almost negligible expression of AXL **(Fig. 2C)**. We further tested AXL localisation by immunostaining in MDAMB231, A549 and CaLu1 cells **(Fig 2E)**. Stable adherent MDAMB231 and A549 cells, where the Golgi is intact, show a strong localisation of AXL with the cis-Golgi marker (GM130), cis-medial Golgi marker (ManII-GFP) and trans-Golgi marker (GalTase-RFP). In CaLu1 cells, the dispersed cis-Golgi, cis-medial and trans-Golgi shows partial overlap with AXL **(Fig 2E)**. Together, they strongly support the presence of AXL at the Golgi and a possible role in its organisation.

### AXL mediated regulation of Golgi organisation in breast cancer cells

MDAMB231, with significant AXL expression, and MCF7, with almost no AXL, make an ideal setting to test the role of AXL in Golgi organisation. Inhibition of AXL using increasing concentrations of R428 (Bemcentinib), a selective ATP competitive inhibitor of AXL, causes both cis- and trans-Golgi to be distinctly disorganised in stable adherent MDAMB231 cells **(Fig 3A**, **Fig 3B)**. AXL is known to regulate Akt activation (Holland et al., 2010), detected by a change in phosphorylation at the Ser473 residue. R428 mediated AXL inhibition is seen to cause a drop in Akt activation across all the concentrations **(Fig 3C)**. This we find is accompanied by a concentration-dependent increase in AXL phosphorylation (Y702) levels on 12 hrs of R428 treatment **(Fig 3D)**. Time kinetics of 1 µM R428 treatment shows a gradual increase in AXL phosphorylation (Y702) levels that is significant at 60 minutes of treatment **(Supp Fig 3A)**. A time course treatment of MDAMB231 cells with R428 (10 min – 60 min) shows a time dependent increase in Golgi disorganization **(Supp Fig. 3C)**. pAkt levels drop and stay low across these treatment timepoints **(Supp Fig 3B)**. While some reports show R428 to decrease AXL phosphorylation at sites – Tyr702(Iida et al., 2017), Tyr779 (Ghosh et al., 2011) and Tyr821 (Holland et al., 2010), another study using R428 treatment shows variable effect on AXL phosphorylation (Chen et al., 2018). Chen et. al. has reported R428 to cause an initial immediate drop in AXL phosphorylation (Tyr702), followed by an increase with time (over 6 hours)(Chen et al., 2018) To address this ambiguity we also tested the pTyr779 AXL antibody in our studies, but found its detection to be very poor (data not shown). In AXL lacking MCF7 cells, R428 treatment does not affect Golgi **(Fig 3E)** or pAkt **(Fig 3F)** levels, further confirming its effect on Akt activation and Golgi organisation in MDAMB231 to be AXL-dependent. The phospho-AXL (Y702) band detected in MDAMB231 cells, is also lost in MCF7 cell lysates confirming its specificity **(Supp Fig. 3D)**. Going ahead we have hence looked at loss of Akt activation and increase in phosphorylation of AXL (Y702) as outcomes of AXL inhibition in these cells.

**Figure 3.**
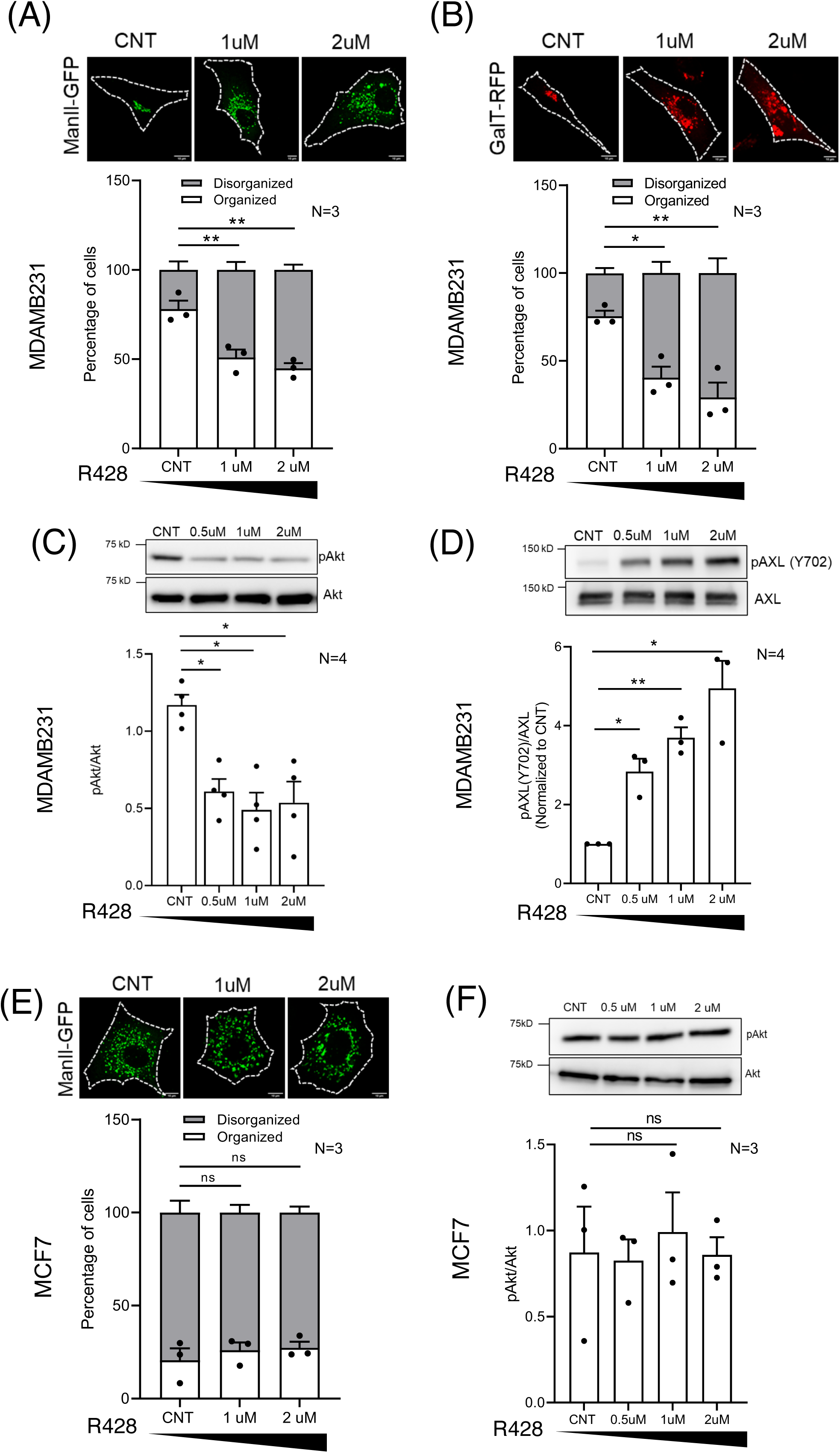
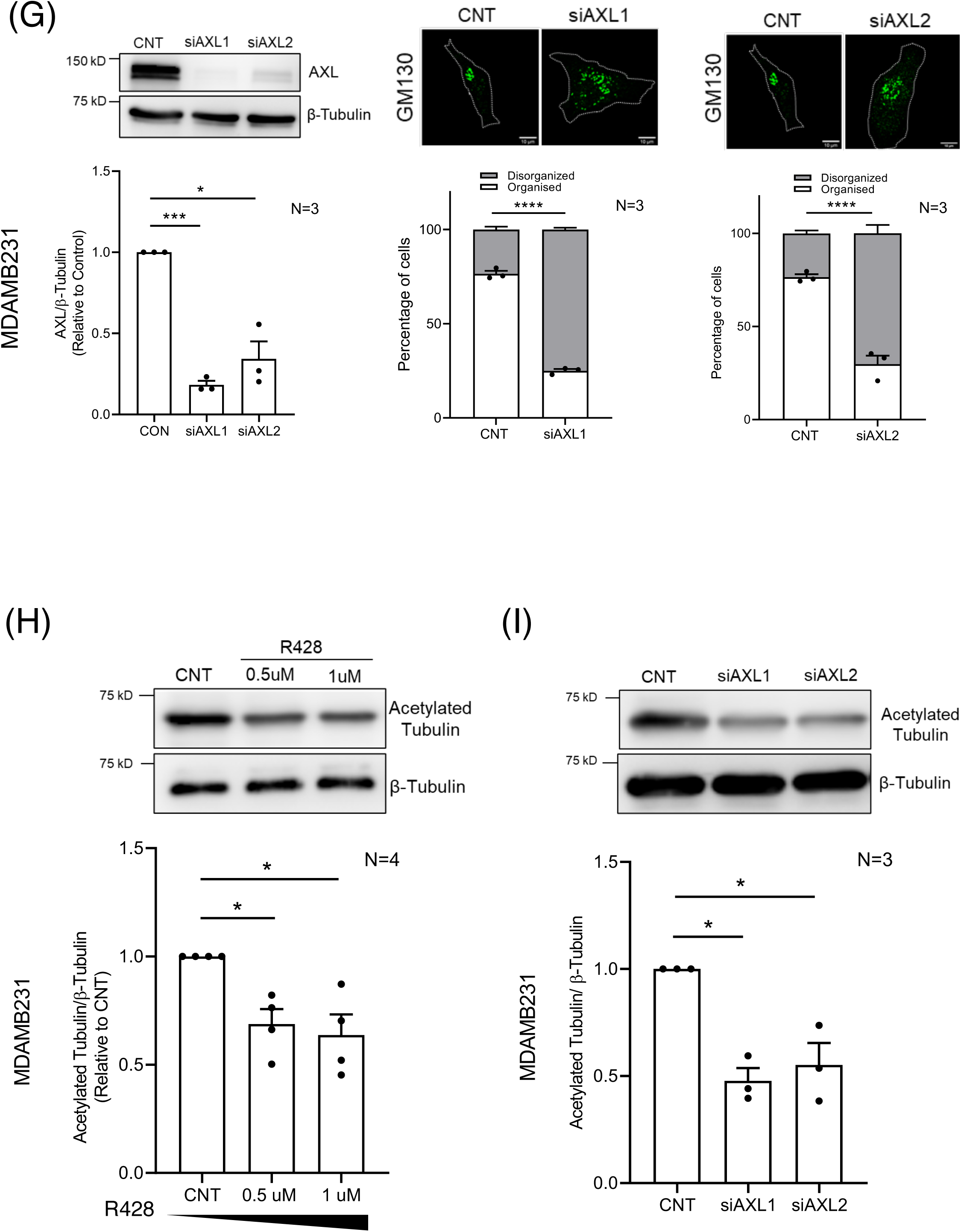
AXL mediated regulation of Golgi organisation in breast cancer cells. **(A, B)** Representative cross-sectional images of the predominant Golgi phenotype in MDAMB231 cells expressing **(A)** ManII-GFP (green) or **(B)** GalTase-RFP (red), treated with DMSO (CNT), or R428 (1μM or 2μM) are shown. Percentage distribution profile for cells (n ≥150) shows organized (white) and disorganized (grey) Golgi in adherent cells across treatments. The graphs represent mean±SEM of percentage distribution profile from three independent experiments. The black bar below the graph represents the gradient of increasing concentration of R428 treatment. **(C, D)** Representative western blots for **(C)** S473 phosphorylated Akt (pAkt) and total Akt (Akt) and **(D)** Y702 phosphorylated AXL (pAXL) and total AXL (AXL) in cell lysates from DMSO (CNT), 0.5μM, 1μM and 2μM R428 treated MDAMB231 cells. Graph represents ratio of densitometric band intensities as mean±SEM from four independent experiments. The black bar below the graph represents the gradient of increasing concentration of R428 treatment. **(E)** Percentage distribution profile for MCF7 cells (n ≥150) expressing cis-medial Golgi marker ManII-GFP with organized (white) and disorganized (grey) Golgi, in DMSO (CNT), 1μM and 2μM R428 treated cells. Representative cross-sectional confocal images of the predominant Golgi phenotype shown. The graph represents mean±SEM of percentage distribution from three independent experiments. The black bar below the graph represents the gradient of increasing concentration of R428 treatment. **(F)** Representative western blots for S473 phosphorylated Akt (pAkt) and total Akt (Akt) in cell lysates from DMSO (CNT), 0.5μM, 1μM and 2μM R428 treated MCF7 cells. Graph represents ratio of densitometric band intensities as mean±SEM from three independent experiments. The black bar below the graph represents the gradient of increasing concentration of R428 treatment. **(G)** Representative western blots for AXL in cell lysates from control (CNT), siAXL1 and siAXL2 treated MDAMB231 cells. Graph represents ratio of densitometric band intensities as mean±SEM from three independent experiments. Representative cross-sectional images of the predominant Golgi phenotype in GM130 (green) immunostained CNT, siAXL1 and siAXL2 MDAMB231 cells. Percentage distribution profile of cells (n ≥150) showing organized (white) and disorganized (grey) Golgi in adherent control (CNT), siAXL1 and siAXL2 treated MDAMB231 cells. The graphs represent mean±SEM of percentage distribution from three independent experiments. Representative western blots for acetylated tubulin and β-tubulin in cell lysates from **(H)** DMSO (CNT), 0.5μM, 1μM R428 treated and **(I)** CNT, siAXL1 and siAXL2 treated MDAMB231 cells. Graphs represent ratio of densitometric band intensities as mean±SEM from four and three independent experiments normalized to control respectively. The black bar below the graph represents the gradient of increasing concentration of R428 treatment. Statistical analysis was done using one way ANOVA multiple comparisons test with Tukey’s method for error correction, for the distribution profiles, Mann-Whitney U test for non-normalised and single sample t-test for normalised (with respect to control) western blotting results respectively. All scale bars shown are 10 µm. (*p≤0.05, **p ≤ 0.01, ***p≤ 0.001, ****p ≤ 0.0001, ns= not significant).

siRNA-mediated knockdown of AXL, using two previously verified individual siRNA sequences (Holland et al., 2010), cause a significant and comparable reduction in AXL levels and disorganisation of the Golgi in MDAMB231 cells **(Fig 3G)**. Loss of AXL is further seen to cause a loss in detection of phospho-AXL (Y702) levels confirming its specificity **(Supp Fig 3D)**. Loss of AXL upon knockdown also caused a drop in Akt activation confirming AXL levels and activation both to be vital in regulating downstream Akt activation **(Supp Fig 3E)**. We further tested if AXL inhibition and knockdown mediated disruption of Golgi organization can affect Golgi function. The Golgi apparatus is known to be a major hub for non-centrosome microtubules to nucleate from (Chabin-Brion et al., 2001). Nucleation and stabilization of the Golgi microtubules is extensively regulated by distinct post-translational modifications such as acetylation (Eshun-Wilson et al., 2019) and could be affected by Golgi organisation (Brodsky et al., 2022; Sanders & Kaverina, 2015). Stable microtubules detected by their acetylation showed a significant drop on R428 treatment **(Fig. 3H)** and siRNA mediated AXL knockdown **(Fig 3I),** reflecting their observed change in Golgi organisation **(Fig 3A, 3B, 3G)**. Together this suggests AXL expression and activation contribute to the regulation of Golgi organisation, which could impact Golgi function.

### AXL-Arf1 crosstalk regulates Golgi organisation in adherent MDAMB231 cells

We further tested if R428 mediated AXL inhibition affects AXL localisation at the Golgi. In MDAMB231, R428 treatment causes AXL to be displaced from the Golgi, reflected in reduction of its co-localisation with GM130 stained cis-Golgi **(Fig 4A)**. No significant change in GM130 levels were seen on R428 treatment **(Supp Fig 4A)**. Earlier studies including ours have shown active Arf1 localisation at the Golgi to be crucial for regulating its organisation and function (B.R. et al., 2023; Ward et al., 2001). The levels and activation status of Arf1 are known to be deregulated in cancers (Casalou et al., 2016, 2020; Xie et al., 2016). Haines et.al have shown in MDAMB231 cells Arf1 can bind AXL in immunoprecipitation studies (Haines et al.), supporting the possible role their crosstalk at the Golgi could have to regulate Golgi organisation. On R428 treatment of MDAMB231 cells, total Arf1 levels while unaffected **(Fig 4B),** showed a small but significant decrease in active Arf1 levels **(Fig 4C)**. GGA3-GST pulldown of Active Arf1 was also seen to bring down AXL with it; this association not affected by R428 treatment **(Fig 4D)**. On AXL knockdown with both tested siRNAs, a significant drop in Arf1 activation was seen **(Fig. 4E)**. This suggests both AXL levels and activation to be critical for Arf1 activation and consequently regulating Golgi organization. R428 did not affect levels of Arf GEF GBF1 **(Supp. Fig 4C)**, known to regulate adhesion-dependent Arf1 activation. Its localization at the Golgi could still contribute to the differential regulation of Arf1, though the GBF1 antibody did not support immunodetection in our hands. We were able to use the ABD-GFP construct to detect active Arf1 localization in cells (B.R. et al., 2023). We find AXL and active Arf1 to colocalise at the Golgi in adherent MDAMB231, which is lost upon R428 treatment **(Fig 4F)**. Loss of ABD-GFP (active Arf1) from the Golgi (GM130) on R428 treatment **(Supp Fig 4B)**, could reflect the loss in Arf1 activation status (seen in pulldown studies) **(Fig 4C)** which could drive Golgi disorganisation. Confirming this regulation, we find overexpressed constitutively active Q71L-Arf1, but not wild-type Arf1, to rescue Golgi organisation in R428 treated cells **(Fig 4G)**. Additionally, targeting Arf1 activation using Arf GEF, GBF1 targeting inhibitor, Golgicide A (GCA) causes the Golgi to be disorganised in adherent MDAMB231 cells **(Supp Fig 4D)**, accompanied by an increase in phospho-AXL (Y702) levels (no change in total AXL levels), possibly reflecting its inhibition as well **(Fig 4H & I)**. This also reflected in loss of AXL localised to the Golgi **(Fig 4J)**. Together, this data lends much credibility to the role AXL-Arf1 crosstalk and their regulation of each other’s activation and localization at the Golgi could have in controlling Golgi organisation.

**Figure 4.**
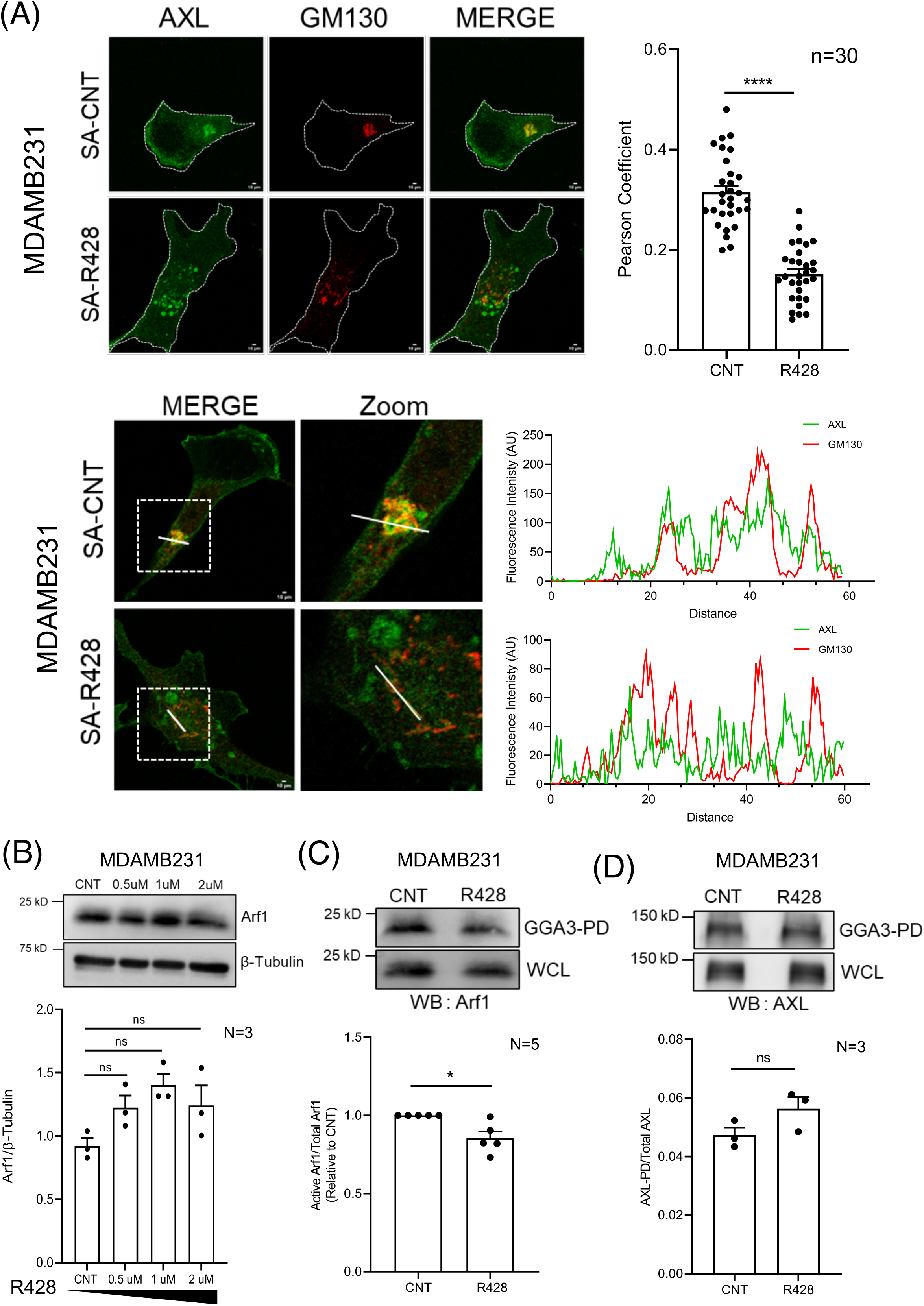
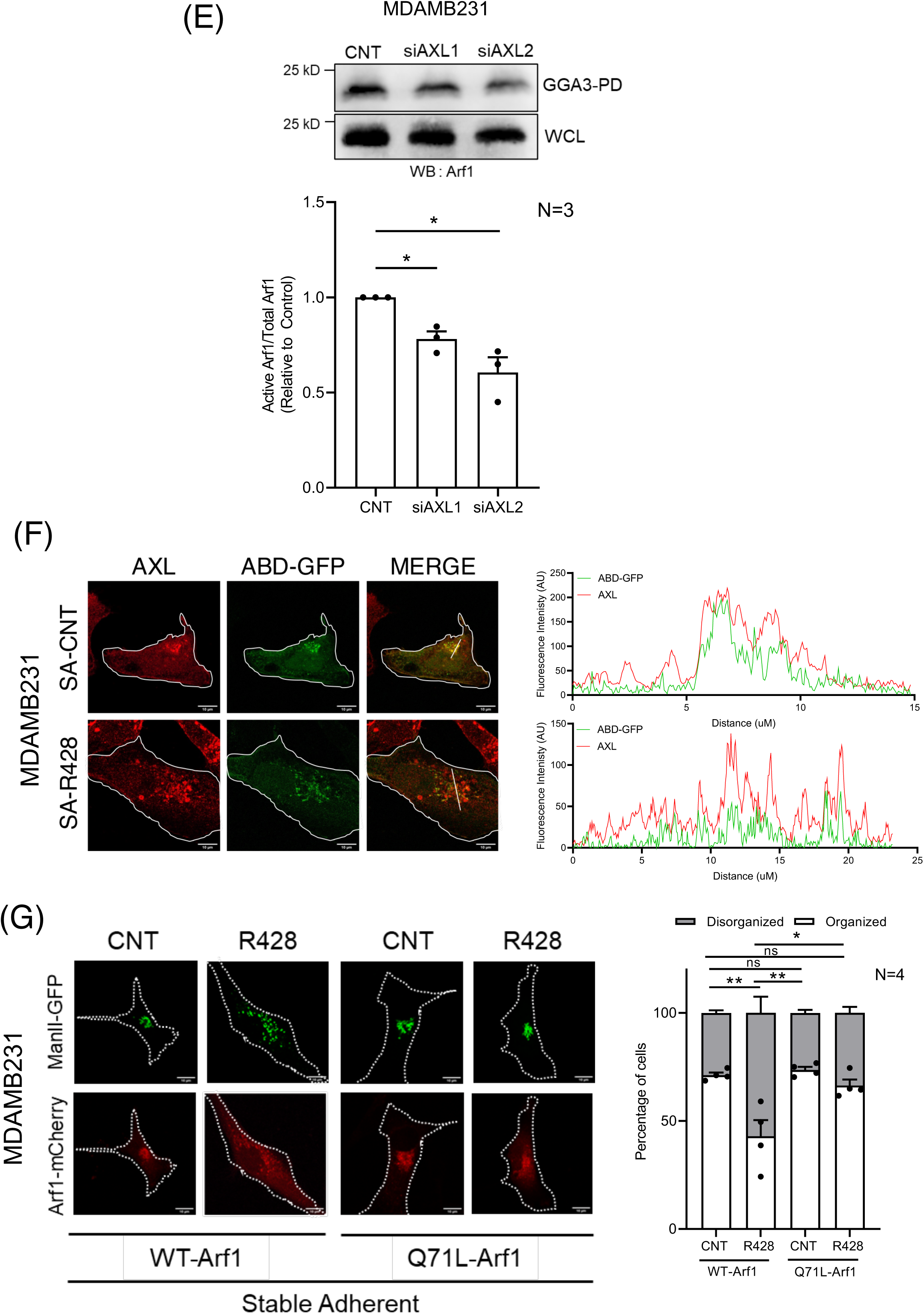
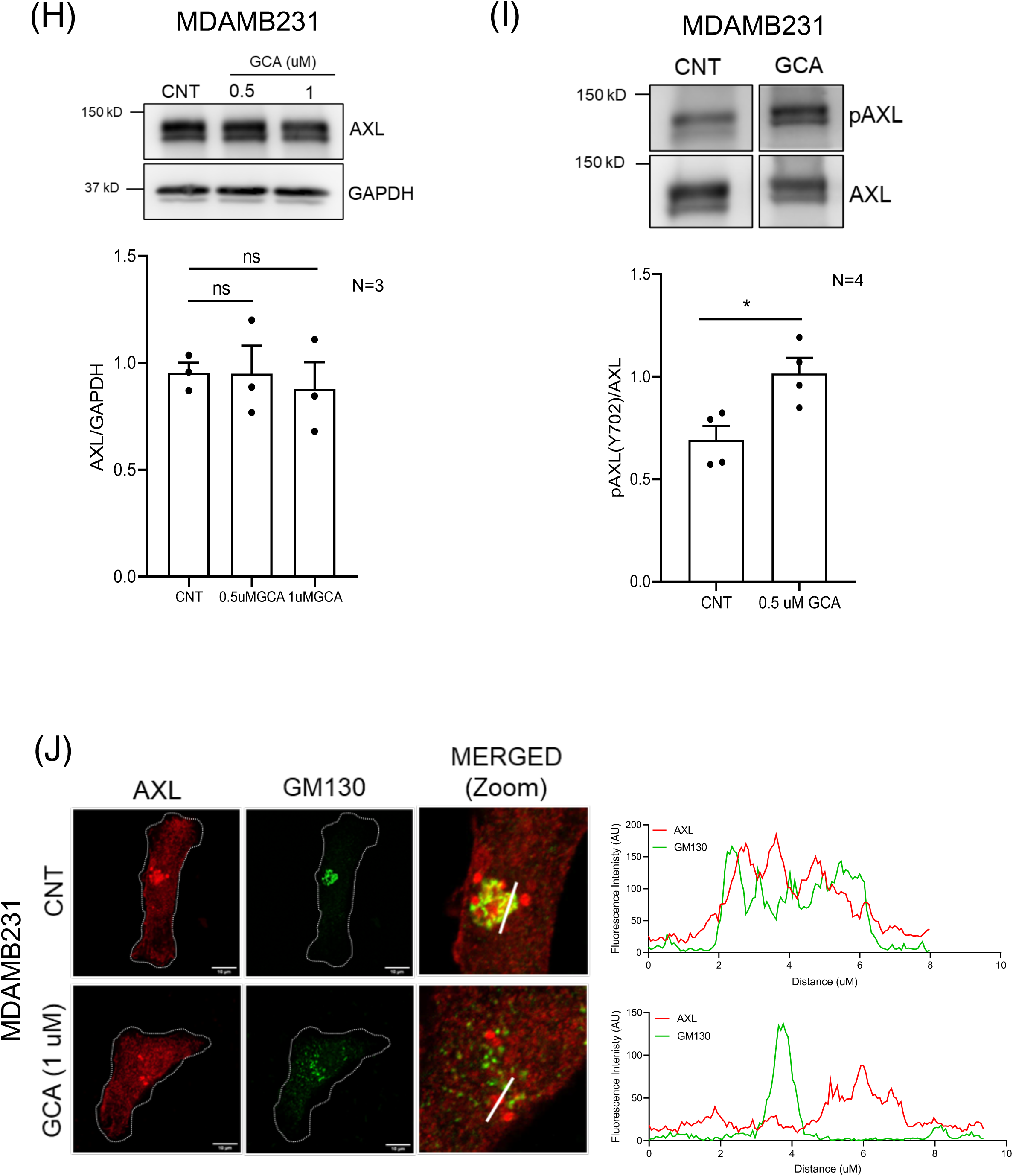
AXL-Arf1 crosstalk regulates Golgi organisation in adherent MDAMB231 cells. **(A) Top Panel** - Representative deconvoluted Z-stack maximum intensity projection (MIP) images for AXL (green) and GM130 (red) immunostained stable adherent DMSO (SA-CNT) or R428 (SA-R428) treated MDAMB231 cells. Individual channel and merged images shown. The graph shows the mean±SEM of Pearson’s correlation coefficient for colocalization of AXL and GM130 (n=30 cells) from three independent experiments. **(A) Lower Panel** – Representative cross-section merged image along with corresponding zoomed image (marked box). Line plot analysis shows fluorescence intensity profile for AXL (green) and GM130 (red) along a solid line in merged MDAMB231 cell image treated with DMSO (SA-CNT) and R428 (SA-R428). **(B)** Representative western blots for Arf1 and β-tubulin in cell lysates from DMSO (CNT), 0.5μM, 1μM and 2μM R428 treated MDAMB231 cells. Graph represents ratio of densitometric band intensities as mean±SEM from three independent experiments. The black bar below the graph represents the gradient of increasing concentration of R428 treatment. Representative western blots for **(C)** Arf1 (WB: Arf1) **(D)** AXL (WB: AXL) in pull down using GST-GGA3 (active Arf1) and in whole-cell lysate (WCL) of DMSO (CNT) and R428 treated MDAMB231 cells. Graph represents ratio of densitometric band intensities in GGA3-PD to WCL as mean±SEM from (C) five (normalized to control) and (D) three independent experiments. **(E)** Representative western blots for Arf1 (WB: Arf1) in pull down using GST-GGA3 (active Arf1) and in whole-cell lysate (WCL) of control (CNT) and AXL knockdown (using siAXL1 and siAXL2) MDAMB231 cells. Graph represents ratio of densitometric band intensities in GGA3-PD to WCL as mean±SEM from three independent experiments, normalized to control. **(F)** Representative cross-sectional images and line plot analysis for stable adherent (SA) MDAMB231 cells expressing ABD-GFP (green) and immunostained for AXL (red), treated with DMSO (SA-CNT) or R428 (SA-R428). Individual channel and merged images shown. Line plot profile for lines marked are shown next to their respective images. **(G)** Representative cross-sectional images of stable adherent MDAMB231 cells expressing constitutively active Arf1 (Q71L-Arf1-mCherry) (red) or wild type-Arf1 (WT-Arf1-mCherry) (red) and cis-medial Golgi marker - ManII-GFP (green) shows the predominant Golgi organization phenotype in DMSO (CNT) and R428 treated cells. Graph shows percentage distribution profile for control and R428 treated WT-Arf1 and Q71L-Arf1 expressing cells (n ≥100) with organized (white) or disorganized (grey) Golgi. The graph represents mean±SEM of percentage distribution from four independent experiments. **(H)** Representative western blots for AXL and GAPDH in cell lysates from DMSO (CNT), 0.5μM and 1μM Golgicide A (GCA) treated MDAMB231 cells. Graph represents ratio of densitometric band intensities as mean±SEM from three independent experiments. **(I)** Representative western blots for Y702 phosphorylated AXL (pAXL) and total AXL in cell lysates from DMSO (CNT), 0.5μM Golgicide A (GCA) treated MDAMB231 cells. Graph represents ratio of densitometric band intensities as mean±SEM from four independent experiments. **(J)** Representative cross-sectional images and line plot analysis for colocalization of AXL (red) and GM130 (green) immunostained adherent MDAMB231 cells, treated with DMSO (CNT) and 1μM GCA for 30 mins. Individual channel and zoomed merged images shown. Line plot profile for lines marked are shown next to their respective images. Statistical analysis was done using one way ANOVA for Pearson’s colocalization analysis and distribution profile. Mann-Whitney U test was used for non-normalised and single sample t test for normalised (with respect to control) western blotting results respectively. All scale bars shown are 10 µm. (*p≤0.05, **p ≤ 0.01, ***p ≤ 0.001, ****p ≤ 0.0001, ns= not significant).

### Role of AXL-Arf1 crosstalk in loss of adhesion mediated Golgi disorganisation in MDAMB231 cells

Adhesion-dependent Golgi organisation in MDAMB231 mimics the ’normal’ phenotype in cells known to be regulated by adhesion-dependent Arf1 activation (Singh et al., 2018). This leads us to ask if AXL-Arf1 crosstalk could also be regulated by adhesion to affect Golgi organisation. In MDAMB231 cells, loss of adhesion causes a significant drop in active Arf1 levels **(Fig 5A)**. This does not affect levels of AXL bound to GGA3-GST beads bound to active Arf1 in stable adherent and suspended cells **(Fig 5B)**. Loss of adhesion we find causes a significant increase in AXL phosphorylation (Y702) **(Fig 5C)** which as discussed earlier could reflect its inhibition. This increase in phospho-AXL (Y702) is seen at early (10 mins) and late (120 mins) times of suspension (**Supp Fig 5A)**. Loss of adhesion is also accompanied by a distinct loss in Akt activation (drop in pAkt levels) **(Fig 5D)**, seen at early (10 min) and late (120 min) times of suspension **(Supp Fig. 5B)**.

**Figure 5.**
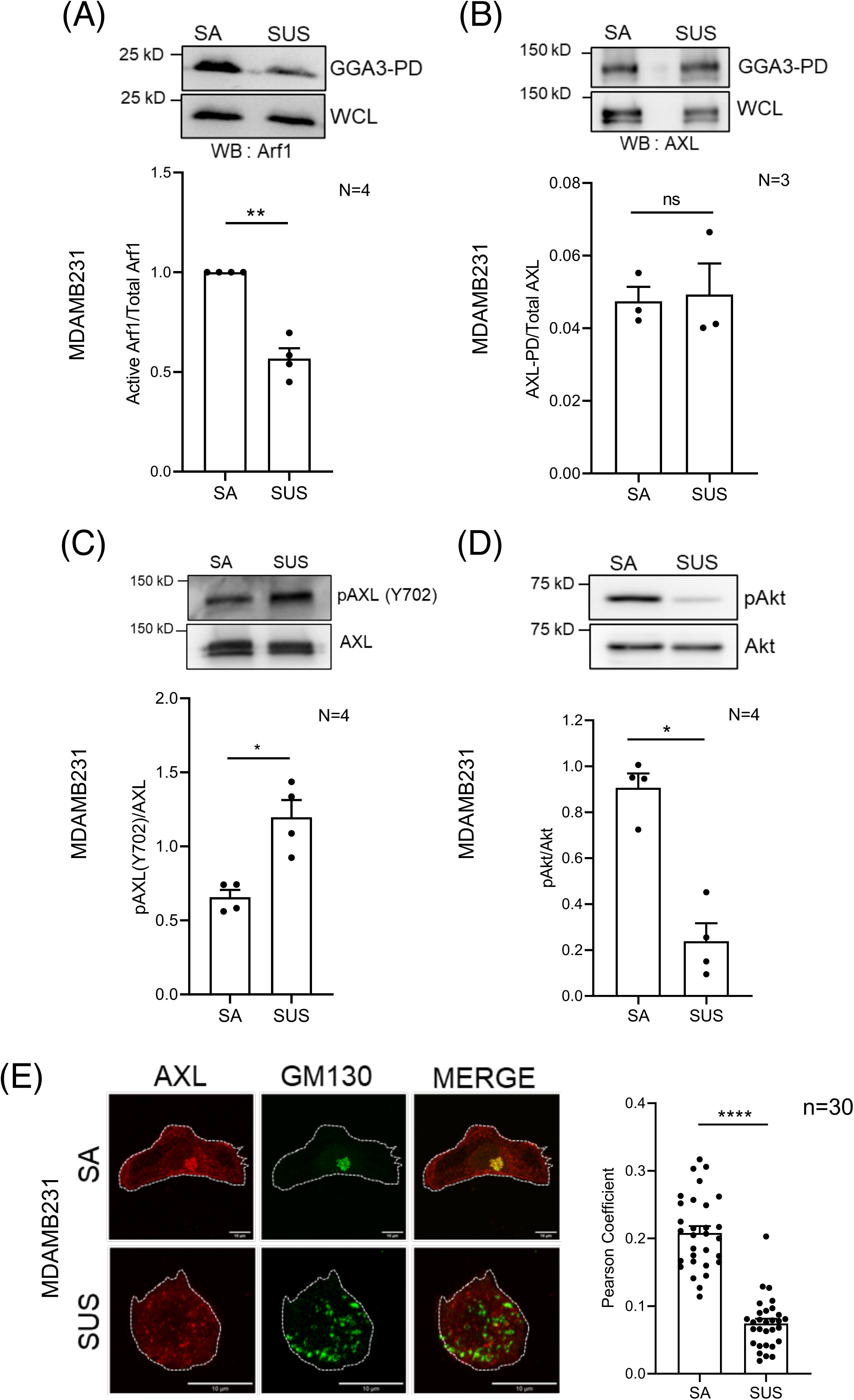
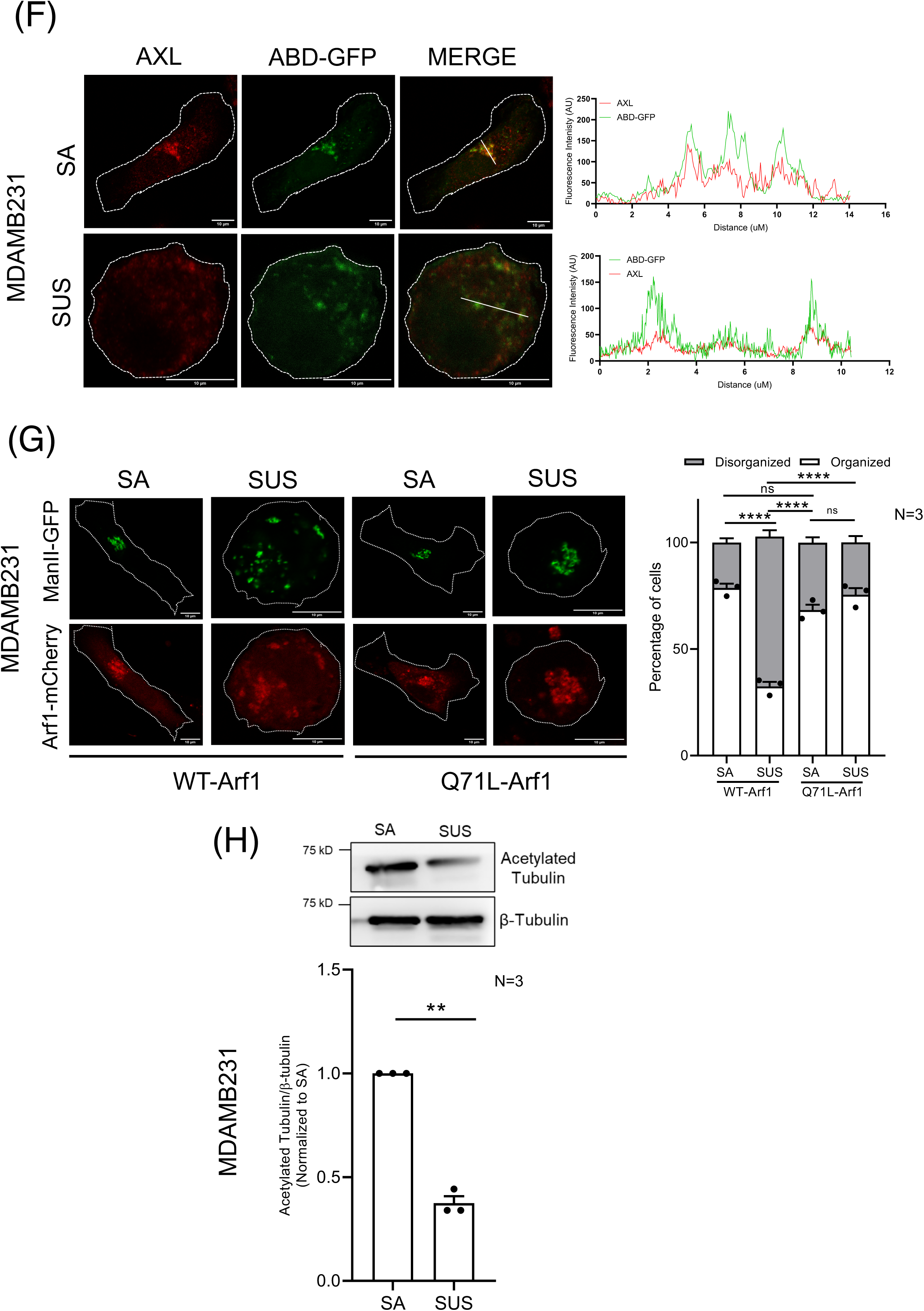
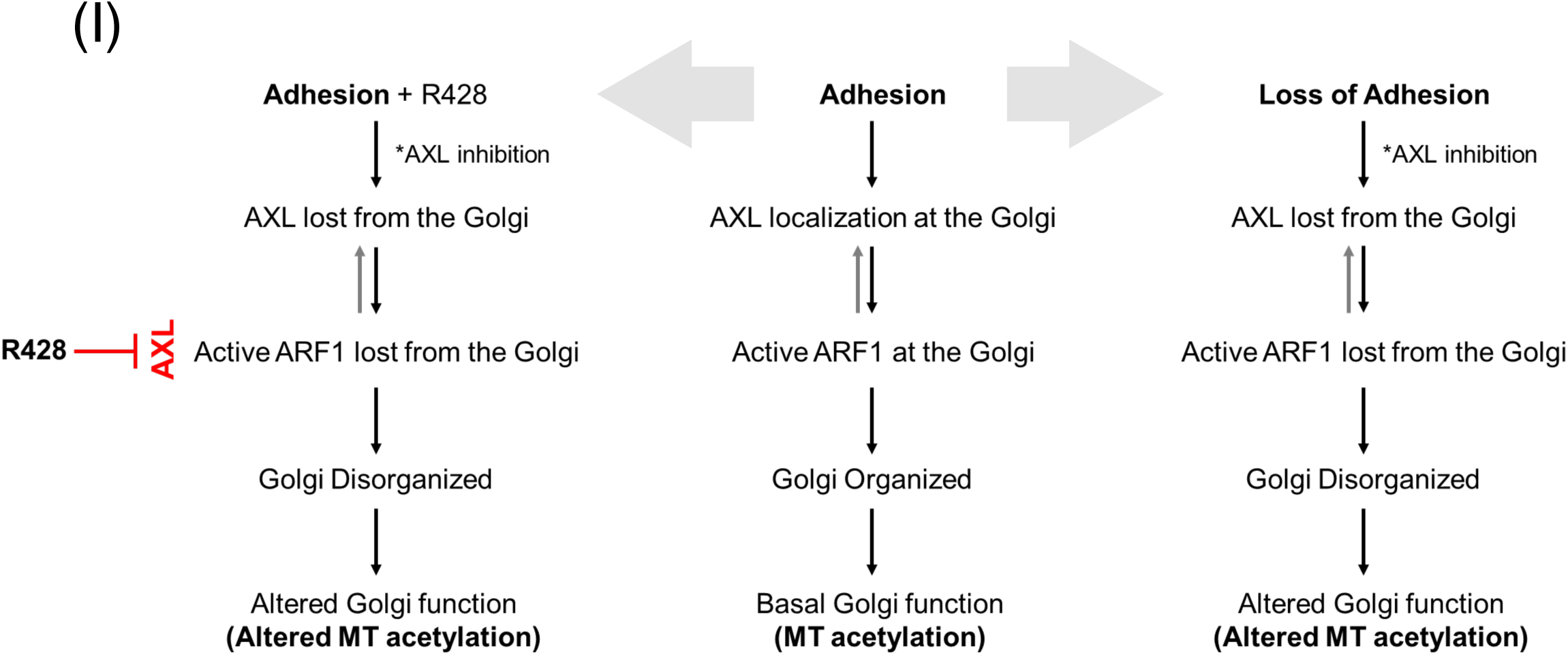
Adhesion-dependent regulation of AXL and Arf1. Role in loss of adhesion mediated Golgi disorganisation in MDAMB231 cells. **(A, B)** Representative western blots for **(A)** Arf1 (WB: Arf1) **(B)** AXL (WB: AXL) in pull down using GST-GGA3 (GGA3-PD) and in whole-cell lysate (WCL) of stable adherent (SA) and non-adherent (SUS) MDAMB231 cells. Graph represents ratio of densitometric band intensities in GGA3-PD to WCL as mean±SEM from (A) four (normalized to control) and (B) three independent experiments. Representative western blots for **(C)** Y702 phosphorylated AXL (pAXL-Y702) and AXL **(D)** S473 phosphorylated Akt (pAkt) and Akt in cell lysates of stable adherent (SA) and non-adherent (SUS) MDAMB231 cells. Graph represents ratio of densitometric band intensities as mean±SEM from (C, D) four independent experiments. **(E)** Representative deconvoluted maximum intensity projection (MIP) images show immunostained AXL (red) and cis-Golgi GM130 (green), in stable adherent (SA) and non-adherent (SUS) MDAMB231 cells. The graph shows the mean±SEM of Pearson’s correlation coefficient for colocalization of AXL and GM130 (n=30 cells) from three independent experiments. **(F)** Representative cross-sectional images and line plot analysis for stable adherent (SA) and non-adherent (SUS) MDAMB231 cells expressing ABD-GFP (green) and immunostained for AXL (red). Individual channel and merged images shown. Line plot profile for lines marked are shown next to their respective images. **(G)** Representative images shown for stable adherent (SA) and non-adherent (SUS) MDAMB231 cells expressing constitutively active Arf1 (Q71L-Arf1-mCherry) (red) or wild type-Arf1 (WT-Arf1-mCherry) (red) and cis-medial Golgi marker ManII-GFP (green). Graph represents the percentage distribution profile of WT-Arf1 and Q71L-Arf1 expressing cells (n ≥150 cells) with organized (white) or disorganized (grey) Golgi. The graph represents mean±SEM from three independent experiments. **(H)** Representative western blots for acetylated tubulin and β-tubulin in cell lysates of stable adherent (SA) and non-adherent (SUS) MDAMB231 cells. Graph represents ratio of densitometric band intensities as mean±SEM from three independent experiments, normalized to SA. **(I)** Schematic describes the AXL-Arf1 crosstalk and its regulation of Golgi organization and function downstream of adhesion, adhesion upon R428 treatment (Adhesion+R428) and loss-of adhesion in MDAMB231 cells. It highlights the comparable outcomes R428 treatment has to loss of adhesion. Statistical analysis was done using one-way ANOVA multiple comparisons test with Tukey’s method for error correction, for Pearson’s colocalization analysis and distribution profile. Mann-Whitney U test was used for non-normalised western blotting results and single sample t test for normalised (with respect to control) western blotting results. All scale bars shown are 10 µm. (*p≤0.05, **p ≤ 0.01, ***p≤ 0.001, ****p≤ 0.0001, ns= not significant).

Furthermore, localisation of AXL **(Fig 5E)** as well as Active Arf1 (detected by ABD-GFP) **(Fig 5F)** seen at the organised Golgi in adherent cells is lost as the Golgi disorganises upon loss of adhesion **(Supp Fig 5C)**. GBF1 levels were also seen to be unaffected on loss of adhesion **(Supp. Fig 5D)**. In MDAMB231 cells overexpression of constitutively active Q71L-Arf1 is also seen to prevent the loss of adhesion-mediated Golgi disorganisation **(Fig 5G)**. Loss of adhesion mediated Golgi disorganization similarly caused a drop in microtubule acetylation **(Fig. 5H)** as observed upon R428 treatment and AXL knockdown **(Fig 3H, 3I)**. Hence, in these cells, loss of adhesion mimics the effect R428 mediated regulation of AXL has on Golgi organisation in adherent cells. These results also reveal how such cells could indeed use the AXL-Arf1 crosstalk to regulate the Golgi in physiological circumstances.

### AXL regulates adhesion-dependent Golgi organisation in lung cancer cells

Lung cancer A549 and CaLu1 cells have a Golgi phenotype that differs from non-transformed lung epithelial BEAS2B cells **(Supp Fig 1A)**. Adherent CaLu1 cells have a disorganised Golgi (Fig 1A) and A549 cells an organised Golgi, that is retained on loss of adhesion **(Fig 1D)**. R428 mediated inhibition of AXL causes a dose-dependent drop in phospho-Akt levels (S473) **(Fig 6C and 6D)** and an increase in phospho-AXL (Y702) levels **(Fig 6E and 6F)** in both cell lines. R428 mediated AXL inhibition in A549 cells causes the Golgi to disorganise **(Fig 6A)**, but does not affect the inherently disorganised Golgi in CaLu1 cells **(Fig 6B**). Interestingly, lower R428 concentrations (1μM) caused significant disorganisation of cis-(GM130), cis-medial (ManII-GFP), and trans-Golgi (GalTase-RFP) in suspended A549 cells **(Fig 6G)**. siRNA-mediated knockdown of AXL in A549 cells done using the siRNA #2 optimised earlier was seen to cause the Golgi to be prominently disorganized in stable adherent and suspended A549 cells **(Fig 6H)**, suggesting AXL can regulate the Golgi in A549 cells but possibly be regulated by adhesion differently **(Fig 6G)**. Both R428 and siRNA mediated AXL knockdown are seen to cause a significant reduction in Akt activation in A549 cells. **(Fig 6I)**.

**Figure 6.**
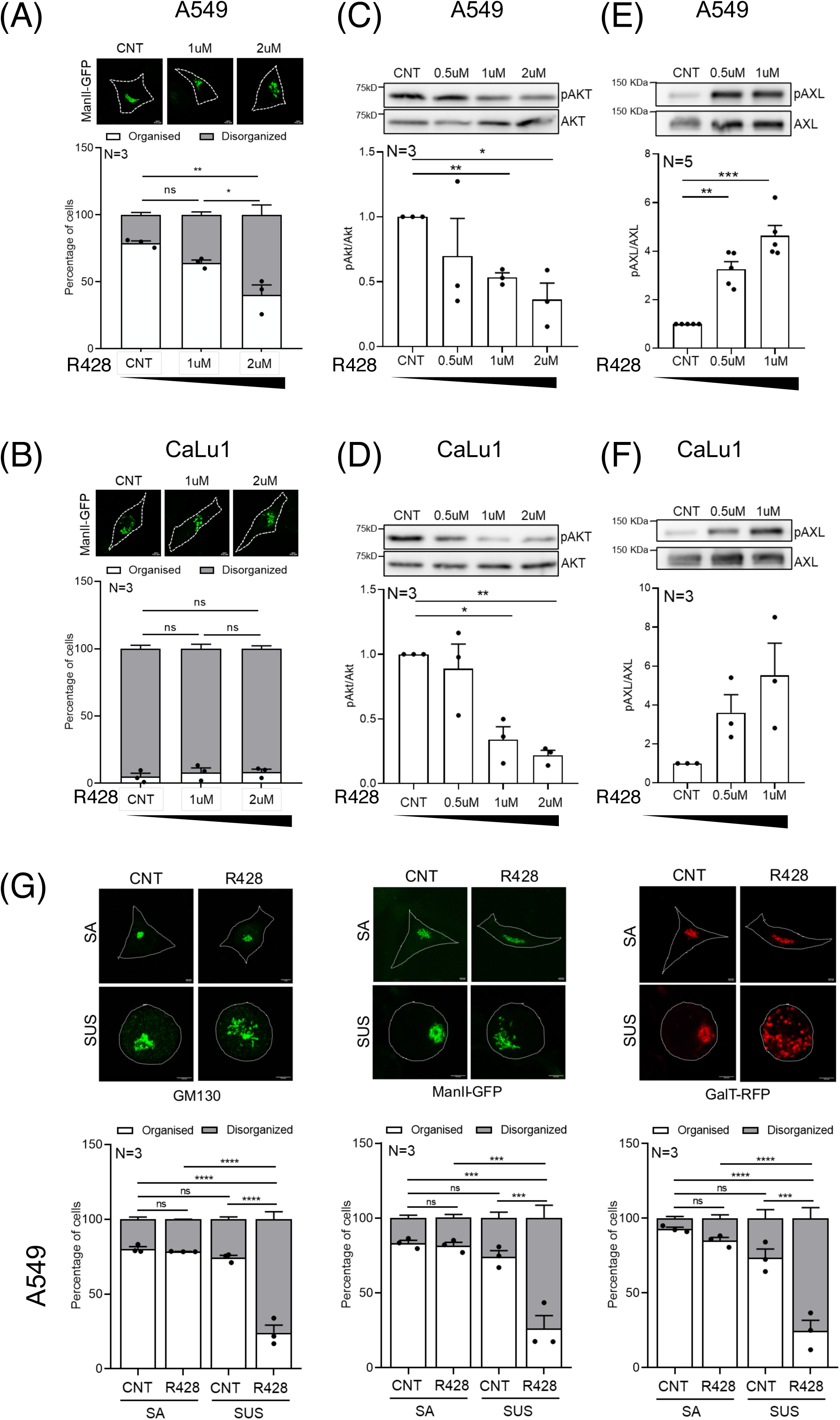
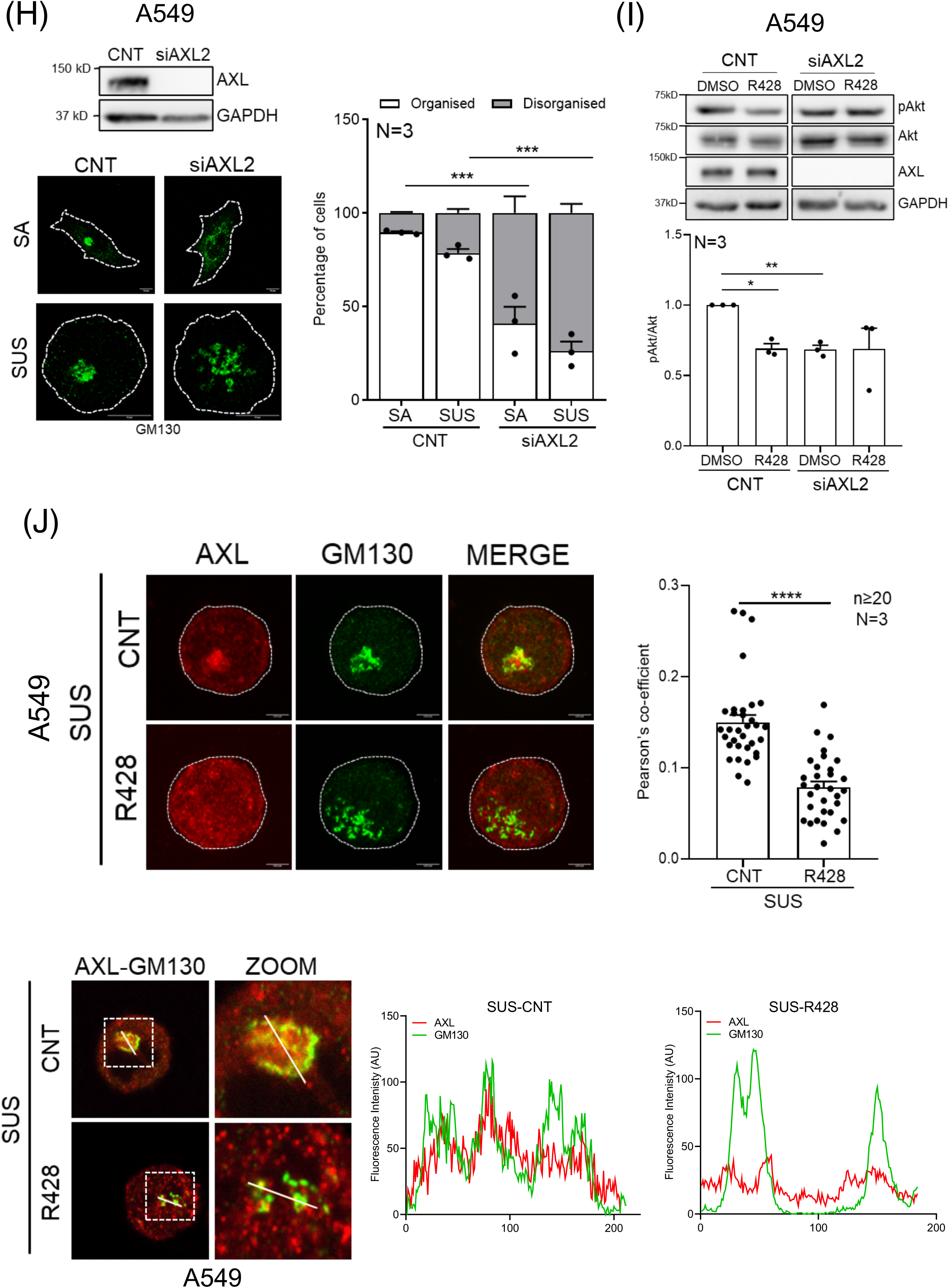
AXL regulates adhesion-independent Golgi organization in lung cancer cells. Representative cross-section images of the predominant Golgi phenotype in **(A)** A549 and **(B)** CaLu1 cells expressing cis-medial Golgi marker-ManII-GFP (green) treated with DMSO (CNT) or R428 (1μM or 2μM) are shown. Percentage distribution profile for cells (n ≥100) shows organized (white) and disorganized (grey) Golgi in adherent cells across treatments. The graphs represent mean±SEM from three independent experiments. The black bar below the graph represents the gradient of increasing concentration of R428 treatment. **(C, D)** Representative western blots for S473 phosphorylated Akt (pAkt) and total Akt (Akt) in cell lysates from DMSO (CNT), 0.5μM, 1μM and 2μM R428 treated **(C)** A549 and **(D)** CaLu1 cells. **(E, F)** Representative western blots for Y702 phosphorylated AXL (pAXL) and total AXL (AXL) in cell lysates from DMSO (CNT), 0.5μM and 1μM R428 treated A549 **(E)** and CaLu1 **(F)** cells. Graphs represent ratio of densitometric band intensities as mean±SEM from three or five independent experiments as indicated in graphs. The black bar below the graph represents the gradient of increasing concentration of R428 treatment. **(G)** Percentage distribution profile for A549 cells (n ≥150) immunostained for cis-Golgi marker -GM130 (green), expressing cis-medial Golgi marker -ManII-GFP (green) or expressing trans-Golgi marker -GalTase-RFP (red) with organized (white) and disorganized (grey) Golgi, in DMSO (CNT) or R428 treated stable adherent (SA) or non-adherent (SUS) cells. Representative deconvoluted z-stack maximum intensity projection (MIP) images of the predominant Golgi phenotype are shown. The graph represents mean±SEM of percentage distribution from three independent experiments. **(H)** Representative deconvoluted maximum intensity projection (MIP) images show the predominant Golgi organization phenotype, immunostained for cis-Golgi marker GM130 (green) in in control (CNT) and AXL knockdown (siAXL2 treated) stable adherent (SA) vs non-adherent (SUS). Representative western blots for AXL and GAPDH to validate AXL knockdown in cell lysates from CNT vs siAXL2 treated A549 cells. Graph represents mean±SEM of percentage distribution profile for cells (n ≥200 cells) with organized (white) and disorganized (grey) Golgi from three independent experiments. **(I)** Representative western blot shows the detection of S473 phosphorylated Akt (pAkt) and total Akt (Akt) and total AXL (AXL) and GAPDH in cell lysates from DMSO (CNT) and R428 treated control (CNT) and AXL knockdown (siAXL2) A549 cells. The graphs represent ratio of densitometric band intensities as mean±SEM from three independent experiments. **(J)** Representative images shown are deconvoluted z-stack maximum intensity projection (MIP), of A549 cells immunostained for AXL (red) and GM130 (green), in non-adherent (SUS) cells treated with DMSO (CNT) or R428. The graph shows the mean±SEM of Pearson’s correlation coefficients for colocalization of AXL and GM130 (n*≥*20 cells) from three independent experiments. Shown below are cross-section images along with corresponding zoomed image of boxed region marked by dotted line. Line plot shows fluorescence intensity profile along the line for AXL (red) and GM130 stained cis-Golgi (green) in DMSO (CNT) or R428. Statistical analysis was done using one-way ANOVA multiple comparisons test with Tukey’s method for error correction, for the distribution profiles, single sample t test for normalised (with respect to control) western blotting results and Mann-Whitney U test for Pearson’s colocalization analysis. All scale bars shown are 10 µm. (*p≤0.05, **p ≤ 0.01, ***p ≤ 0.001, ****p ≤ 0.0001, ns= not significant).

We hence tested, AXL localisation at the Golgi (GM130) in non-adherent A549 cells and found it to be reduced significantly on R428 treatment where the Golgi is disorganized **(Fig 6J)**. This suggests that changes in AXL activation and possible crosstalk with Golgi regulators on loss of adhesion might affect Golgi organisation in lung cancer cells. If the AXL-Arf1 cross-talk, seen in MDAMB231, has a role in lung cancer cells becomes of direct interest.

### AXL-Arf1 crosstalk regulates Golgi organisation and function in non-adherent A549 cells

We hence first tested if Arf1 is regulated by adhesion in A549 cells. Active Arf1 levels (GGA3 pulldown) drop significantly on loss of adhesion in A549 cells **(Fig 7A)**. However, this drop is not sufficient to cause the Golgi to disorganise in these cells. In suspended cells, R428 treatment did not affect Arf1 activation **(Fig 7B)** or its binding to AXL in pulldowns **(Fig 7C)**. Loss of adhesion does not affect AXL phosphorylation (Y702) **(Fig 7D),** but on treatment with R428, it increases significantly **(Fig 7E)**, accompanied by a distinct drop in AKT activation (pS473 AKT) **(Fig 7F)**. This when viewed in light of the fact that AXL inhibition (by R428), causes it to leave the Golgi **(Fig 6J),** suggests displacement of Arf1 from the Golgi that could contribute to this regulation. Despite a drop in active Arf11 levels on loss of adhesion, residual active Arf1 seen localized at the Golgi **(Fig 7G)** could allow for its organization to be retained in A549 cells. R428 mediated AXL inhibition causes active Arf1, detected using ABD-RFP, to be significantly displaced from the Golgi driving its disorganization **(Fig 7G)**.

**Figure 7.**
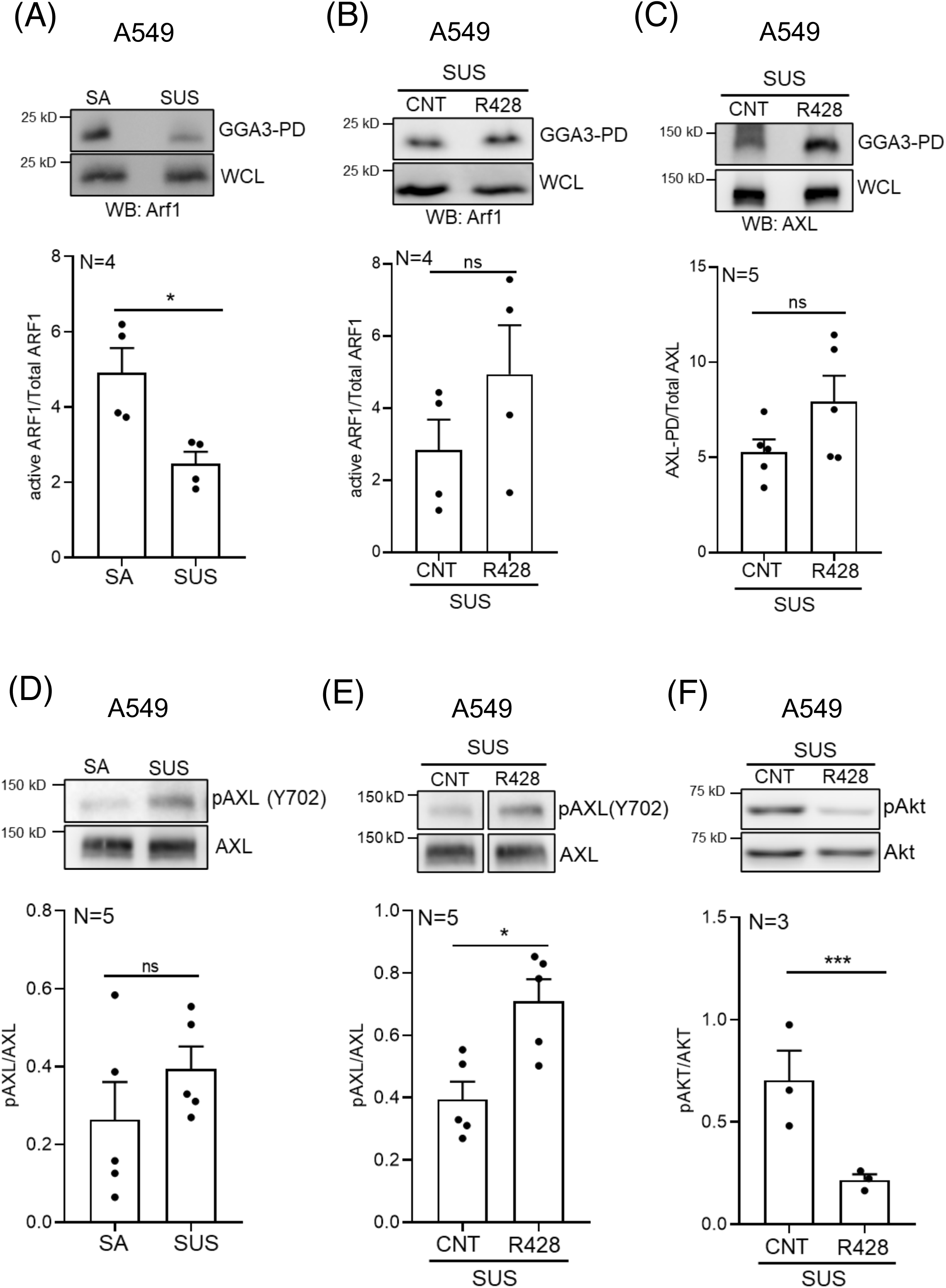
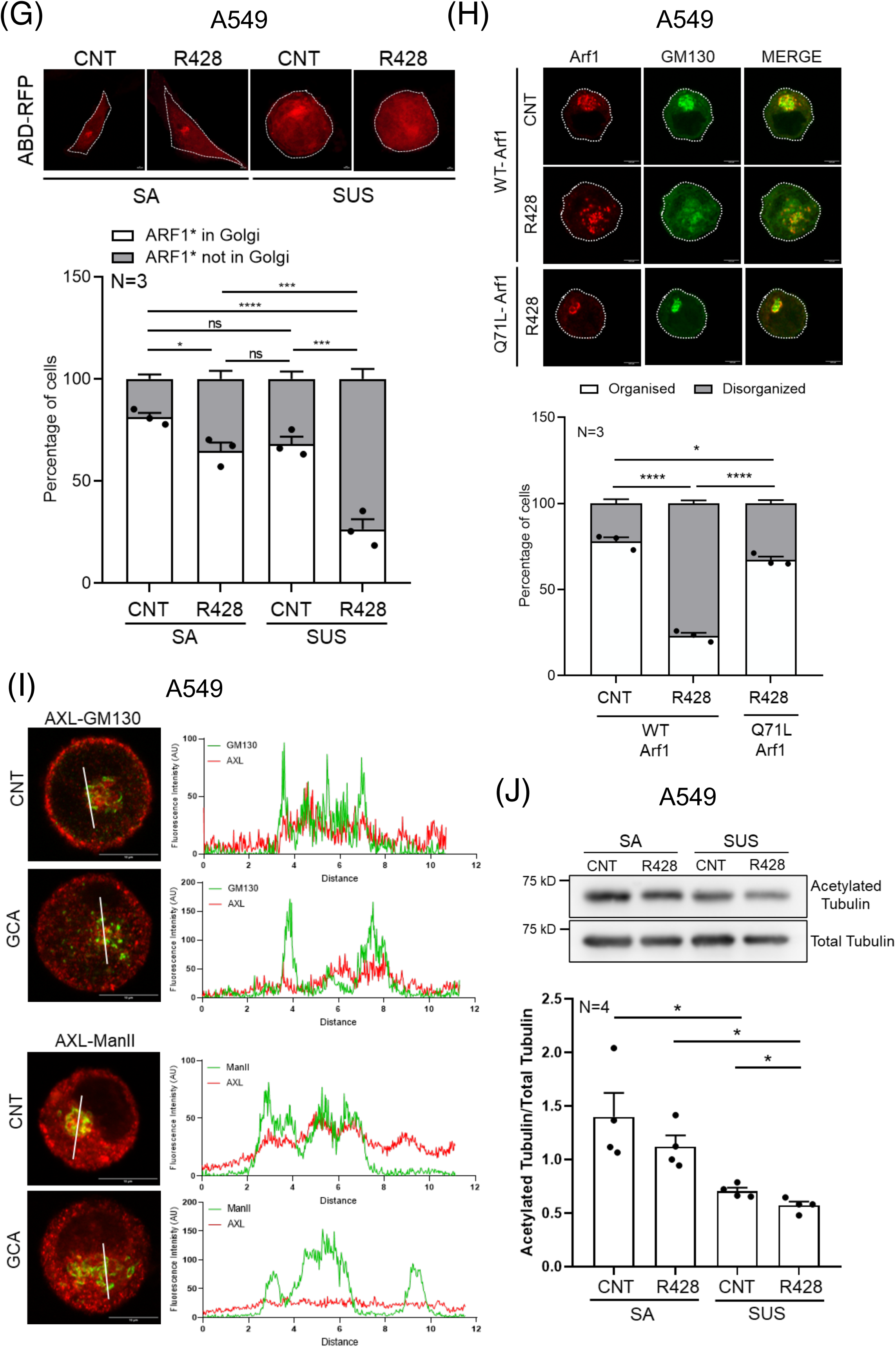
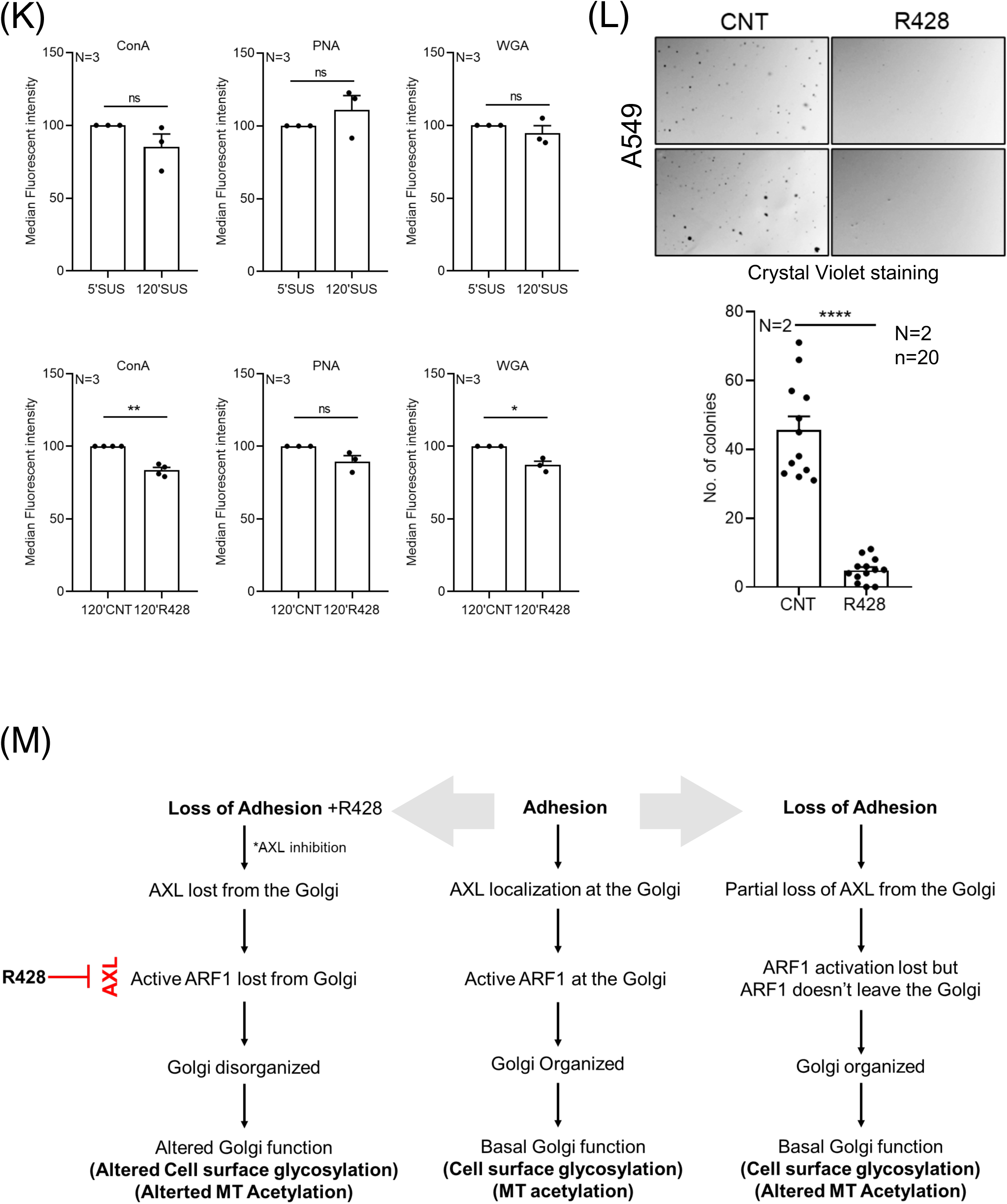
AXL-Arf1 crosstalk regulates Golgi organization in non-adherent A549 cells. Representative western blots for **(A, B)** Arf1 (WB: Arf1) and **(C)** AXL (WB: AXL) in pull down using GST-GGA3 (GGA3-PD) and whole-cell lysate (WCL) of **(A)** stable adherent (SA) and non-adherent (SUS) and **(B, C)** non-adherent (SUS) - DMSO (CNT) and R428 treated A549 cells. Graph represents ratio of densitometric band intensities in GGA3-PD to WCL as mean±SEM from **(A, B)** four and **(C)** five independent experiments. Representative western blots for **(D, E)** Y702 phosphorylated AXL (pAXL), AXL and **(F)** S473 phosphorylated Akt (pAkt) and Akt in cell lysates of **(D)** stable adherent (SA) vs non-adherent (SUS) and **(E, F)** DMSO (CNT) vs R428 treated non-adherent A549 cells. Graph represents ratio of densitometric band intensities as mean±SEM from five or three independent experiments as mentioned in the graphs. **(G)** Percentage distribution profile for cells (n ≥200 cells) expressing ABD-RFP (red) to detect active Arf1 enriched at an intracellular location, confirmed previously to overlap (white) or not overlap with the Golgi marker (grey) in adherent (SA) and non-adherent (SUS) A549 cells treated with DMSO (CNT) or R428. Representative images are cross-section confocal images showing the predominant Golgi phenotype. The graph represents mean±SEM from three independent experiments. **(H)** Representative deconvoluted z-stack maximum intensity projection (MIP) images for A549 cells expressing constitutively active Arf1 (Q71L-Arf1-mCherry) (red) or wild-type-Arf1 (WT-Arf1-mCherry) (red), and immunostained for cis-Golgi marker GM130 (green) in DMSO (CNT) and R428 treated non-adherent A549 cells. Graph shows percentage distribution profile for non-adherent control and R428 treated WT-Arf1 and Q71L-Arf1 expressing cells (n ≥200) with organized (white) or disorganized (grey) Golgi. The graph represents mean±SEM from three independent experiments. **(I)** Representative cross-sectional confocal images with line plot profile for colocalization of immunostained AXL (red) and GM130 (green) or AXL (red) and ManII-GFP (green) expressed in non-adherent A549 cells treated with DMSO (CNT) and 1μM GCA. **(J)** Representative western blot for acetylated tubulin and total tubulin in cell lysates from DMSO (CNT) and R428 treated - stable adherent (SA) and non-adherent (SUS) A549 cells. The graph represents ratio of densitometric band intensities as mean±SEM from four independent experiments. **(K)** Graphs represent mean±SEM of median fluorescent intensities of cell surface bound ConA, PNA and WGA lectins, detected by flow cytometry analysis, at early (5 min) or late (120 min) suspension timepoints and in DMSO (CNT) and R428 treated A549 cells from three independent experiments. **(L)** Representative images of crystal violet stained colonies in soft agar of DMSO (CNT) and R428 treated A549 cells. The graph represents mean±SEM of number of colonies counted from 20 images obtained from two independent experiments. **(M)** Schematic describes the AXL-Arf1 crosstalk and its regulation of Golgi organization and function downstream of adhesion, loss of adhesion and loss of adhesion on R428 treatment (loss of adhesion+R428) in A549 cells. Statistical analysis was done using one-way ANOVA multiple comparisons test using Tukey’s method for error correction for the distribution profiles, Mann-Whitney U test for non-normalised western blotting results and AIG assay, and single sample t-test for normalised (with respect to early suspension time point and control) flow cytometry results. All scale bars shown are 10 µm. (*p≤0.05, **p≤ 0.01, ***p≤ 0.001, ****p≤ 0.0001, ns= not significant).

So, what changes in suspended A549 cells on R428 treatment that causes the Golgi to be dispersed? R428 treatment does not cause a significant change in Arf1 activation **(Fig 7B)** and AXL-Arf1 binding **(Fig 7C)** in non-adherent A549 cells. This suggests the amount of AXL bound to active Arf1 in these suspended cells is likely unaffected on AXL inhibition. R428 treatment also causes a significant increase in AXL phosphorylation (Y702) **(Fig 7D)**. This is accompanied by a significant decrease in Akt activation **(Fig 6C and 6D)** which we have earlier seen to be associated with AXL inhibition **(Fig 6E and 6F)**. R428 treatment leads to a dramatic loss of active Arf1 localisation (detected using ABD-RFP) from the Golgi only in non-adherent A549 cells **(Fig 7G Supp Fig 7)**. Expression of constitutively active Arf1 (Q71L) known to localise to the Golgi, can prevent R428 mediated Golgi dispersal on loss of adhesion in A549 cells **(Fig 7H),** as seen MDAMB231 cells **(Fig 5G)**. This suggests a loss of AXL accompanied by a significant loss of Arf1 from the Golgi is needed for its disorganisation. Interestingly treatment of suspended A549 with GBF1 inhibitor Golgicide-A (GCA) causes the Golgi to disperse with loss of AXL localization **(Fig 7I)**. We also observe a drop in microtubule Acetylation on loss of adhesion in these cells which is accompanied by a drop in Arf1 activation **(Fig7J)**. This is further enhanced on Golgi disorganisation by R428 treatment (AXL inhibition) **(Fig7J)**. These results suggest that in A549 cells both Arf1 activation (K. H. Zhang et al., 2023) and Golgi organisation could regulate microtubule Acetylation.

Cell surface glycosylation changes, detected by fluorescent lectin labelling, are seen as a sensitive and direct measure of changes in Golgi function (Bekier et al., 2017; X. Zhang & Wang, 2016). Quantitative flow cytometric measurements of lectin labelling (ConA, WGA, PNA) in non-adherent A549 cells show no significant change in their glycosylation signature **(Fig 7K)**. R428 treatment of these cells causes a modest but significant change in cell surface bound ConA (binding mannose) and WGA (binding Sialic acid/N-GlcNAc glycans) lectin levels. No change in PNA (binding Galactose and N-Acetyl Galactosamine (N-GalNAc) lectin levels were seen **(Fig 7K)**. This suggests AXL inhibition mediated Golgi disorganisation could affect Golgi function in A549 cells. As reported in earlier studies (Sun et al., 2022), R428 treatment of A549 cells is also seen to significantly affect their anchorage-independent growth **(Fig 7L)**, which could in part be regulated by its effect on Golgi organisation and glycosylation.

Together, these studies in breast and lung cancer cells (MDAMB231 and A549 cells) identify the AXL–Arf1 crosstalk to be vital in regulating Golgi organization and function. This is mediated by AXL binding of Arf1 regulating their recruitment at the Golgi. Regulation of AXL and Arf1 by adhesion could further contribute to their role in Golgi organisation and function.

## DISCUSSION

As a member of the TAM (Tyro3, AXL, Mer) family of RTKs, AXL is frequently overexpressed in multiple malignancies, including breast cancer, non-small cell lung cancer (NSCLC), and acute myeloid leukemia (AML), its elevated levels often correlated with poor prognosis and reduced survival rates (Auyez et al., 2021; Zhu et al., 2019). The functional versatility of AXL, coupled with its involvement in key signaling pathways, underscores its potential as a therapeutic target in cancers. Changes in the organisation and function of the Golgi, including its regulation of cell-surface glycosylation, can enhance cancer cell metastasis (Rambaruth & Dwek, 2011), immune evasion (Demetriou et al., 2001), survival (Petrosyan et al., 2014; Piyush et al., 2017) and drug resistance (Britain et al., 2018; Lopez Sambrooks et al., 2018; Ohashi et al., 2018), making the Golgi an attractive organelle in cancer therapy. (Bhat et al., 2017; Ohashi et al., 2018; Petrosyan et al., 2014). While the role Golgi fragmentation could have in supporting cancer phenotypes has been speculated on (Makhoul et al., 2019; Petrosyan, 2015), understanding how differential Golgi organisation could affect cancers remains unclear. The differential regulation of the Golgi in breast (MDAMB231 vs MCF7) and lung (A549 vs Calu1) cancers and the identification of differentially expressed genes that could regulate Golgi organisation could add significantly to this understanding. The identification of AXL as a primary differentially expressed gene of interest makes its regulation and role at the Golgi relevant.

Seen to prominently localise at the Golgi, loss of AXL protein (siRNA knockdown) and its inhibition (R428 treatment) are both seen to regulate Golgi organisation in breast cancer and lung cancer cells. Both also regulate Akt activation downstream of AXL in MDAMB231, A549, and CaLu1 cells, but this regulation by R428 is missing in AXL-lacking MCF7 cells, confirming the inhibitor’s specificity. While AXL was suggested to localize at the Golgi (Zajac et al., 2020), its distinct overlap with cis-Golgi, cis-medial-Golgi, and trans-Golgi markers in our study confirms this. This localization of AXL at the Golgi is vital for its organisation, mediated by its regulation of Arf1 activation and active Arf1 localization at the Golgi. Gefitinib promotes Arf1 recruitment to AXL, effectively enhancing Arf1 activity (Haines et al., 2015) at the plasma membrane. Their association at the Golgi (AXL binding active Arf1 in GGA3 pulldown and localizing with ABD-GFP) and a distinct regulatory crosstalk shown in our study makes the AXL-Arf1-Golgi connection vital to the role of AXL. AXL inhibition and knockdown mediated drop in Arf1 activation and resulting disorganisation of the Golgi being restored by constitutively active Arf1 (Q71L) confirms their crosstalk and its role. The fact that Golgicide-A mediated Arf1 inhibition regulates AXL phosphorylation and localization suggests this crosstalk to be bidirectional. AXL and Arf1 hence regulate each other’s presence and role at the Golgi, making this crosstalk particularly relevant. In MDAMB231 on loss of adhesion a drop in Arf1 activation accompanied by an increase in AXL phosphorylation and their displacement from the Golgi further suggests this regulatory crosstalk to play a role under physiological circumstances as well. Thus, our study reports for the first time that cell-matrix adhesion regulates AXL activation and its distinct localization at the Golgi. A comparable functional outcome for AXL knockdown, AXL inhibition and loss of adhesion mediated regulation of AXL on Golgi-dependent function is reflected in how they all similarly affect Golgi-dependent microtubule acetylation, which could, as reported in literature, in turn, affect microtubule stability (Eshun-Wilson et al., 2019). Microtubule acetylation is known to be affected by Arf1 activation (K. H. Zhang et al., 2023) and Golgi organisation (Brodsky et al., 2022; Sanders & Kaverina, 2015), making it a vital functional outcome of AXL-dependent regulation of the Golgi in cancers. Golgi associated MTs have been implicated as “fast tracks” for anterograde trafficking (Hao et al., 2020) and regulators of directed cell migration (Wu et al., 2016), which could both have implications for cancers.

Knowing AXL lacking MCF7 cells to have a dispersed Golgi, the effect restoring AXL expression has on the Golgi in these cells could further establish the role their differential expression could have. Additional studies from the lab evaluating the mechanosensory role of AXL in breast cancer cells, have tested and seen AXL to restore Golgi organisation in MCF7 (Manuscript in bioRxiv). The implications of restored AXL levels on Golgi function hence also becomes relevant (Manuscript in bioRxiv).

Knowing AXL to be differentially expressed in breast cancer and lung cancer cells, a differential regulation of AXL-Arf1-Golgi crosstalk in these cells could add to the relevance this regulatory pathway has. MDAMB231 and A549 cells have an intact Golgi when adherent, and AXL knockdown and inhibition are both seen to affect Golgi organisation, mediated by displacement of AXL from the Golgi. Unlike MDAMB231 cells, in A549 cells, a lower R428 concentration, causes a pronounced disorganisation of the Golgi on loss of adhesion, but does not significantly affect the Golgi in stable adherent cells. Additionally, AXL inhibition led to displacement of active Arf1 along with AXL from the Golgi in both cell lines. However, this affects Arf1 activation only in MDAMB231 cells. This suggests AXL-mediated regulation of Arf1 and the Golgi could also vary across cell types. The binding between AXL and active Arf1 is however conserved, and seen in both MDAMB231 and A549 cells.

In response to loss of adhesion, a drop in AXL and Arf1 activation along with displacement of both AXL and active Arf1 from the Golgi causes Golgi disorganisation in non-adherent MDAMB231 cells. However, in A549 cells we observed that loss of adhesion does not cause Golgi disorganization, deeming the AXL mediated regulation of the Golgi to be adhesion independent. This is further reflected in how AXL activation, localization at the Golgi and active Arf1 localization at the Golgi are all retained in non-adherent A549 cells. In response to loss of adhesion A549 cells show a drop in Arf1 activation, but no change in AXL activation (Y702 phosphorylation). Further, in A549 suspended cells, both AXL and active Arf1 remain at the Golgi, making their regulation distinct from MDAMB231. R428 treatment displaces AXL and active Arf1 (ABD-RFP) leading to Golgi disorganization in suspended A549 cells. Like in MDAMB231 cells, constitutively active Arf1 was able to reverse R428 Golgi phenotype in lung cancer cells supporting the presence of an AXL-Arf1-Golgi connect in these cells.

A functional change in the cell surface glycosylation levels for ConA and WGA bound lectins, a distinct change in microtubule acetylation and suppression of anchorage independent growth all suggest AXL targeting and resulting disruption of the Golgi to have function implications for cancer progression. Being able to target only the AXL localised at the Golgi thus becomes of direct interest. Changes in the A549 glycocalyx can further regulate viral infectivity in lung epithelial cells (Lai et al., 2025) that could add to the role AXL-Arf1-Golgi pathway could have in disease.

Studies suggest AXL targeting overcomes drug resistance against chemotherapeutic agents and RTK inhibitors across different cancers such as NSCLC, breast cancer, pancreatic cancer, acute myeloid leukaemia, ovarian cancer, and glioblastoma (Auyez et al., 2021; Debruyne et al., 2016; Hong et al., 2008; Okura et al., 2020; Scaltriti et al., 2016; Z. Zhang et al., 2012). Interestingly, targeting the Golgi organization in NSCLC tumors is shown to promote resistance to EGFR-specific RTK inhibitors (Ohashi et al., 2018), and Arf1 inhibition promotes resistance to Chemotherapy and RTK inhibitors, in breast cancer cells (Haines et al., 2015; Luchsinger et al., 2018). We see that AXL, Arf1 and Golgi are all independently shown to promote drug resistance in cancers, while these players could possibly have a concerted role in the process. Similarly, AXL (Bi et al., 2017; Onken et al., 2016; Zajac et al., 2020), Arf1 (Casalou et al., 2016; Lewis-Saravalli et al., 2013), and Golgi organization (Isaji et al., 2014; Millarte & Farhan, 2012; Zajac et al., 2020) are independently reported to enhance cancer cell migration. Our data suggests the AXL-Arf1-Golgi organization axis, could have a role in both drug resistance and cancer cell migration that remains to be fully explored.

Finally, understanding the regulatory and functional implication of AXL being localised at the Golgi and plasma membrane remains of much interest. Arf1 is well established to have roles at both sites (Boulay et al., 2008; Donaldson et al., 2005; Lewis-Saravalli et al., 2013). This could have implications for the AXL-Arf1 crosstalk at both locations. Does AXL regulate Arf1 at the PM, like we show at the Golgi, remains unclear. When displaced from the Golgi on R428 treatment AXL re-distribution in cells is variable, localised in distinct intracellular structures in MDAMB231 and being more dispersed in A549 cells. Re-localisation of AXL and Arf1 from the Golgi to the PM could have implications for their role at both sites.

While AXL as a transmembrane receptor tyrosine kinase has its ligand binding domain localised outside the cell at the PM (Zhu et al., 2019), access to the same could be limited when localised at the Golgi. Understanding how AXL ligand, Gas6 impacts AXL-Arf1-Golgi organisation could add another aspect to this regulation. An additional role for this pathway could also be in cellular mechanosensing. Shown now to be regulated by adhesion, AXL and ROR2 are reported to alter rigidity sensing by regulating local mechanosensory contractions (Yang et al., 2016). Differential regulation of the Golgi in response to mechanical cues while speculated (Romani et al., 2019), remains to be established. Could the adhesion-AXL-Arf1-Golgi crosstalk define a mechanosensory pathway that could impact cell function is a question our independent studies have also addressed (Manuscript in bioRxiv). Hence, this study and its identification of an AXL-Arf1-Golgi pathway, not just validates the simple screen we began with, but also shows how it could reveal new Golgi regulatory paradigms with implications for normal cell function and disease.

## Acknowledgement

This work is supported by a SERB CRG grant – CRG/2022/001813 to NB. PJ was supported by a fellowship from IISER,Pune. AS is supported by a Prime Minister’s Research Fellowship (PMRF). RM is supported by a fellowship from the Council of Scientific & Industrial Research (CSIR). We acknowledge the extensive support provided by the IISER Pune Microscopy Facility for cellular imaging and the IISER, Pune FACS Facility.

## Author Contributions

The experimental work was led jointly by PJ and AS. Data was recorded, organized and analyzed by PJ, AS, RM and DP. Individual experiments had support from GM, MP and VS. Data was recorded, organized and analyzed by GM, MP and VS. The manuscript was written through the contributions of PJ, AS, RM and NB. All authors have read and given approval to the final version of the manuscript. The authors declare no competing financial interest.

---

**Supp. Figure 1. Adhesion dependent regulation of Golgi organization in non-transformed breast and lung epithelial cells. (A)** BEAS2B cells transfected with cis-medial Golgi marker, ManII-GFP (green) or trans-Golgi marker, GalTase-RFP (red). Representative deconvoluted Z-stacks images for the predominant phenotype shown as maximum intensity projection (MIP) and a zoomed image of the Golgi with surface rendering (SR). Percentage distribution profile of cells (n ≥200) show organized (white) and disorganized (grey) Golgi in stable adherent (SA) and non-adherent (SUS) cells. Graphs represent mean±SEM from three independent experiments. **(B)** Stable-adherent MCF10A cells immunostained for cis-Golgi marker, GM130 (green). Representative cross-sectional confocal images for organized and disorganized Golgi phenotype shown. Percentage distribution profile for cells (n ≥200) show organized (white) and disorganized (grey) Golgi for cells. Graph represents mean±SEM from three independent experiments. **(C)** Representative deconvoluted Z-stacks images for GM130 (green) immunostained in stable adherent (SA), non-adherent (90’ SUSP) and re-adherent (FN 5’) MCF10A cells shown as maximum intensity projection (MIP) and surface rendered (SR) image. Graph shows mean±SEM of discontinuous cis-Golgi (GM130) objects per cell for non-adherent (90’ SUSP) and re-adherent (FN 5’) (n=10 cells). Statistical analysis done using one-way ANOVA multiple comparisons test with Tukey’s method for error correction for the distribution profile, and Mann-Whitney U test for object count analysis. All scale bars shown are 10 µm. (*p≤0.05, **p ≤ 0.01, ***p ≤ 0.001, ****p ≤ 0.0001, ns= not significant).

**Supp Figure 3. AXL mediated regulation of Golgi organisation in adherent MDAMB231 cells.** Representative western blots for **(A)** Y702 phosphorylated AXL (pAXL) and total AXL (AXL) and **(B)** S473 phosphorylated Akt (pAkt) and total Akt (Akt) in cell lysates from DMSO treated for 60 min (DMSO-60 min) and R428 treated for 10, 20, 30, 40, 60 min MDAMB231 cells. Graph represents ratio of densitometric band intensities as mean±SEM from four independent experiments. The black bar below the graph represents the gradient of increasing time of R428 treatment. **(C)** Representative cross-section images of the organised and disorganised Golgi phenotype in GM130 (red) immunostained control (DMSO-12 hrs) or R428 treated for 10, 20, 30, 40, 60 min and 12 hrs MDAMB231 cells. Percentage distribution profile of cells (n ≥150) showing organized (white) and disorganized (grey) Golgi across treatment conditions. The graph represents mean of percentage distribution from one representative experiment. Representative western blots for **(D)** Y702 phosphorylated AXL (pAXL), total AXL (AXL) and GAPDH in cell lysates from control MDAMB231 (MDA-CNT), siAXL1 (AXL KD1), siAXL2 (AXL KD2) and MCF7 cells. **(E)** Representative western blots for S473 phosphorylated Akt (pAkt) and total Akt (Akt) in cell lysates from control (CNT) and siAXL1 treated MDAMB231 cells. Graph represents ratio of densitometric band intensities as mean±SEM from three independent experiments normalized to control. Statistical analysis was done using Mann-Whitney U test for non-normalised and single sample t test for normalised (with respect to control) respectively. All scale bars shown are 10 µm. (*p≤0.05, **p ≤ 0.01, ***p ≤ 0.001, ****p ≤ 0.0001, ns= not significant).

**Supp. Figure 4. AXL-Arf1 crosstalk regulates Golgi organisation in adherent MDAMB231 cells (A)** Representative western blot for GM130 and GAPDH levels in adherent MDAMB231 cells treated with DMSO (CNT) or R428 (0.5μM, 1μM and 2μM). The black bar below the graph represents the gradient of increasing R428 concentration. The graph represents ratio of densitometric band intensities as mean±SEM from three independent experiments. **(B)** Representative cross-section images of cells expressing ABD-GFP (green) and the Golgi immunostained with GM130 (red) in adherent MDAMB231 cells, treated with DMSO (SA-CNT) or R428 (SA-R428). Individual channel and merged images shown. Line plot profile to measure fluorescence intensity for ABD-GFP and GM130 shown next to their respective images. **(C)** Representative western blot for GBF1 and GAPDH levels in adherent MDAMB231 cells treated with DMSO (CNT) or R428 (0.5μM, 1μM, 2μM). The graph represents ratio of densitometric band intensities as mean±SEM from three independent experiments. The black bar below the graph represents the gradient of increasing R428 concentration. **(D)** Percentage distribution profile of cells (n ≥150 cells) showing organized (white) and disorganized (grey) Golgi in adherent MDAMB231 cells, stained with DAPI (blue) and immunostained with GM130 (green) treated with DMSO (CNT) or GCA (0.5μM and 1μM). Representative cross-section confocal images of predominant Golgi organization phenotype shown. The graph represents mean from one experiment. Statistical analysis was done using Mann-Whitney U test for non-normalised western blotting results. All scale bars shown are 10 µm. (*p≤0.05, **p ≤ 0.01, ***p ≤ 0.001, ****p ≤ 0.0001, ns= not significant).

**Supp. Figure 5. Adhesion-dependent regulation of AXL and Arf1. Role in loss of adhesion mediated Golgi disorganisation in MDAMB231 cells. (A-B)** Representative western blots for **(A)** Y702 phosphorylated AXL (pAXL) and total AXL (AXL) and **(B)** S473 phosphorylated Akt (pAkt) and total Akt (Akt) in cell lysates from stable-adherent (SA) and non-adherent MDAMB231 cells suspended for 10, 20, 30, 40, 60 and 120 minutes. The black bar below the graph represents the gradient of increasing time of R428 treatment. The graphs represent ratio of densitometric band intensities as mean±SEM from four independent experiments respectively. **(C)** Representative cross-section images of cells expressing ABD-GFP (green) and the Golgi, immunostained with GM130 (red) in stable adherent (SA) and non-adherent (SUS) MDAMB231 cells. Individual channel and merged images shown. Line plot profile to measure fluorescence intensity for ABD-GFP and GM130 shown next to their respective images. **(D)** Representative western blot for GBF1 and GAPDH in lysates from stable-adherent (SA) and non-adherent (SUS) MDAMB231 cells. The graphs represent ratio of densitometric band intensities as mean±SEM from three independent experiments. Statistical analysis was done using Mann-Whitney U test for non-normalised western blotting results. All scale bars shown are 10 µm. (*p≤0.05, **p ≤ 0.01, ***p ≤ 0.001, ****p ≤ 0.0001, ns= not significant).

**Supp. Figure 7. R428 treatment and its effect on active Arf1 localization at the Golgi (A)** A549 cells expressing ABD-GFP (green) to detect active Arf1 enriched at an intracellular location, seen to overlap with the Golgi immunostained for GM130 (red) in adherent (SA) and non-adherent (SUS) cells treated with DMSO (CNT) or R428. Representative images are cross-section confocal images showing the predominant phenotype. Individual channel and merged images shown. All scale bars shown are 10 µm.

