## Supplementary figures and images for "Receptor tyrosine kinase AXL regulates Golgi organization and function through an adhesion-Arf1 signaling axis in breast and lung cancers"

### Supplementary Figures

#### Supp Figure 1

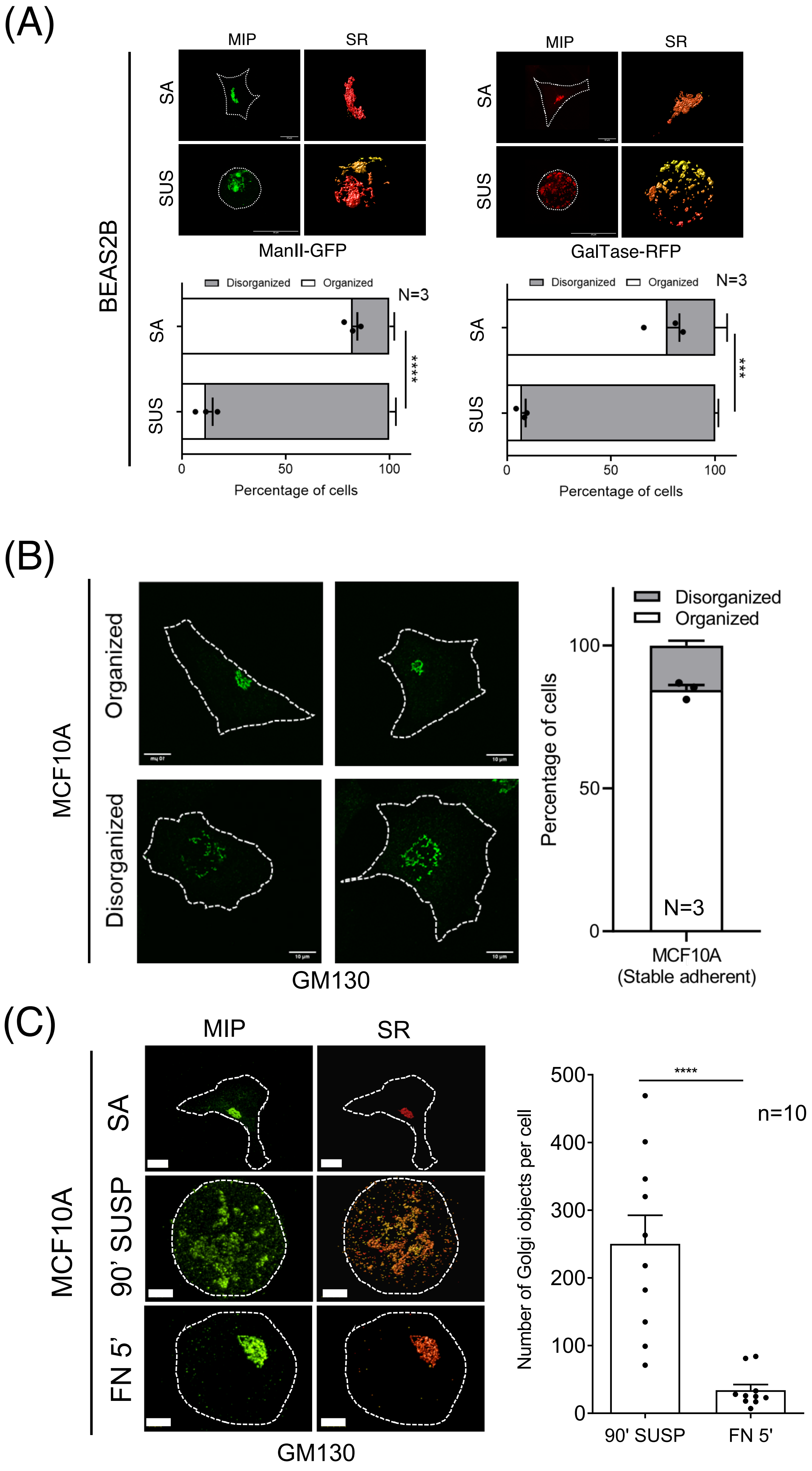

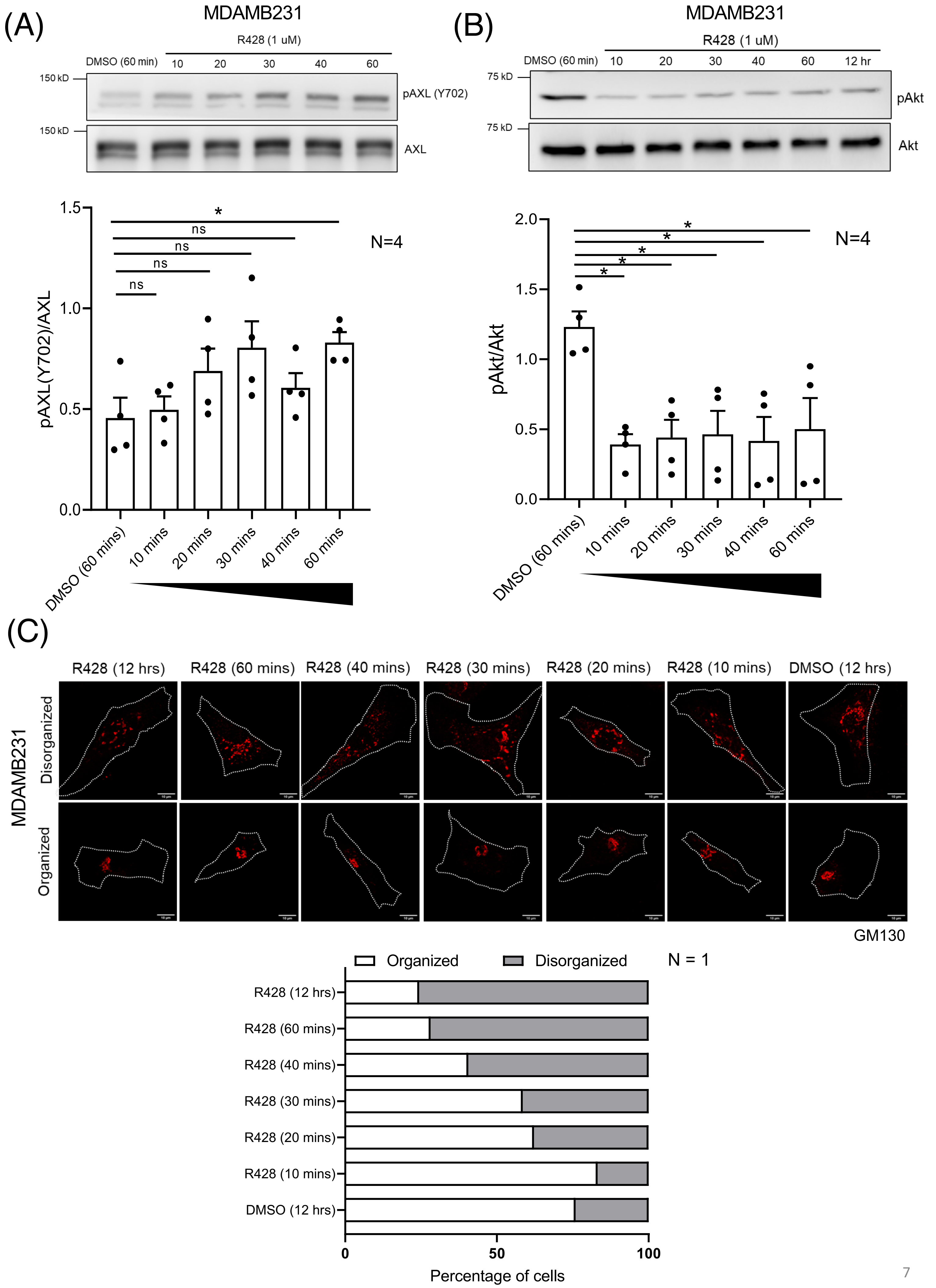

Supp Figure 3

(D)

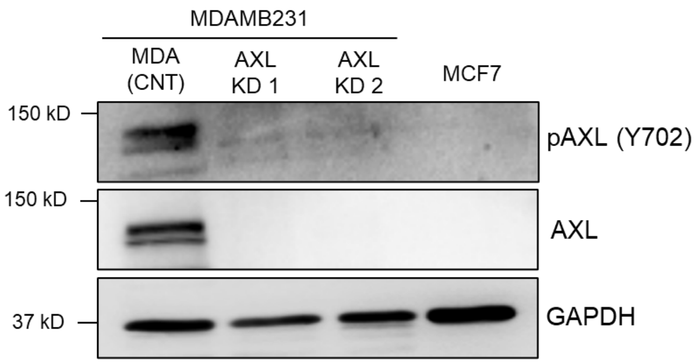


(E)

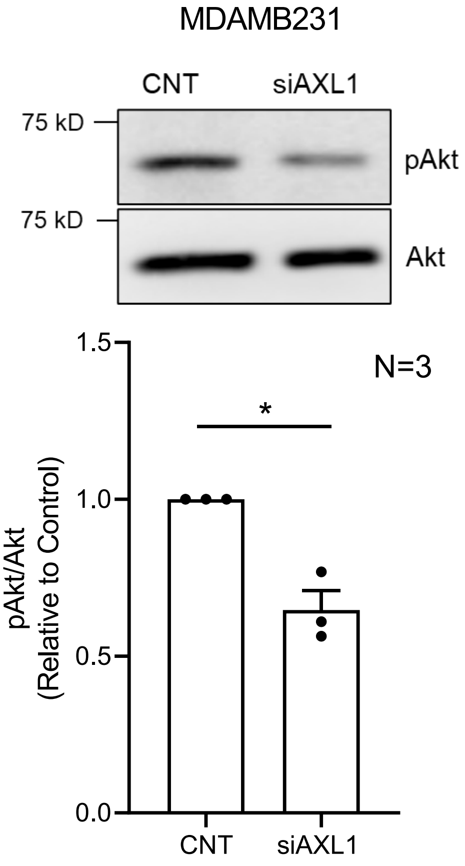

#### Supp Figure 4

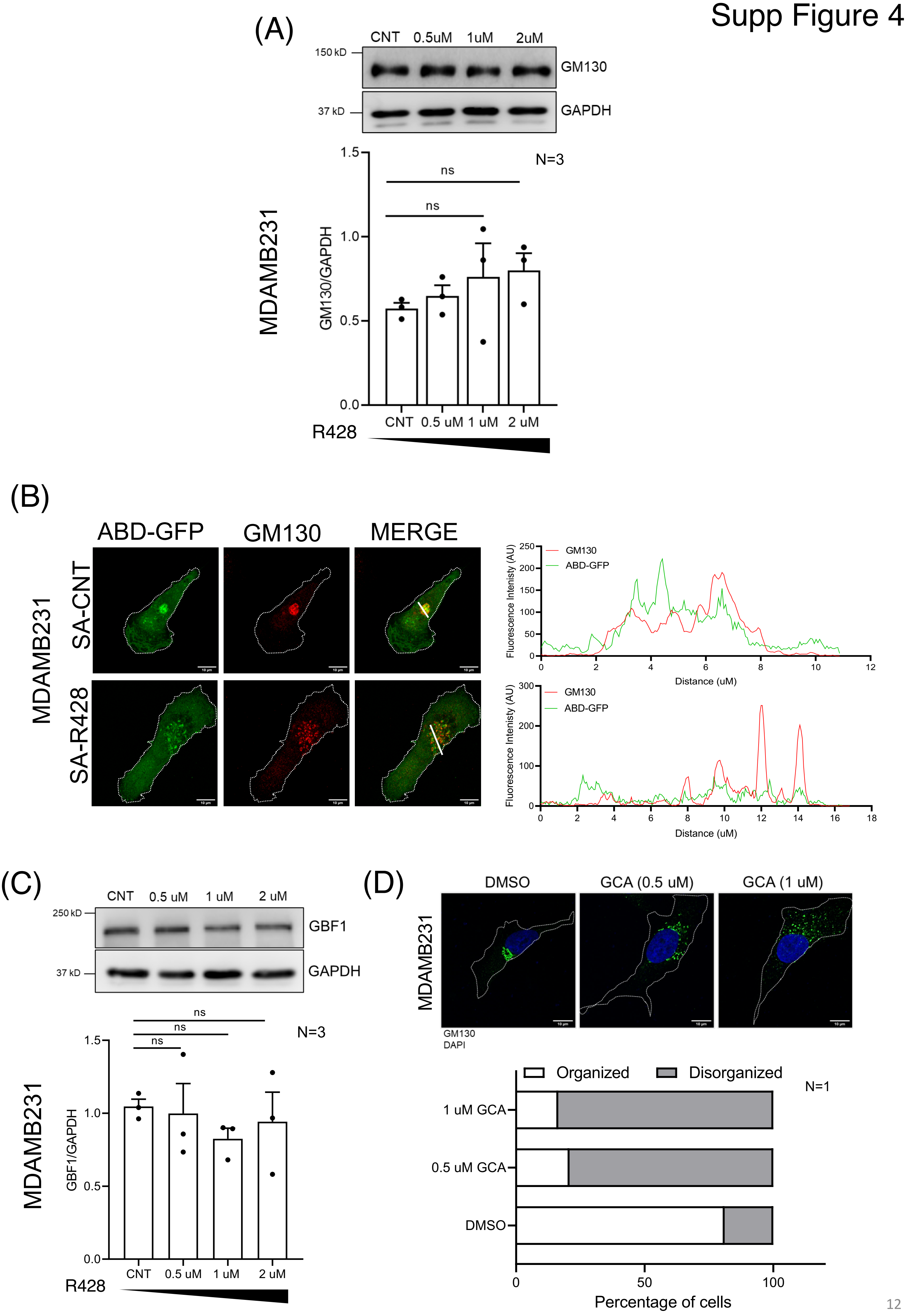

#### Supp Figure 5

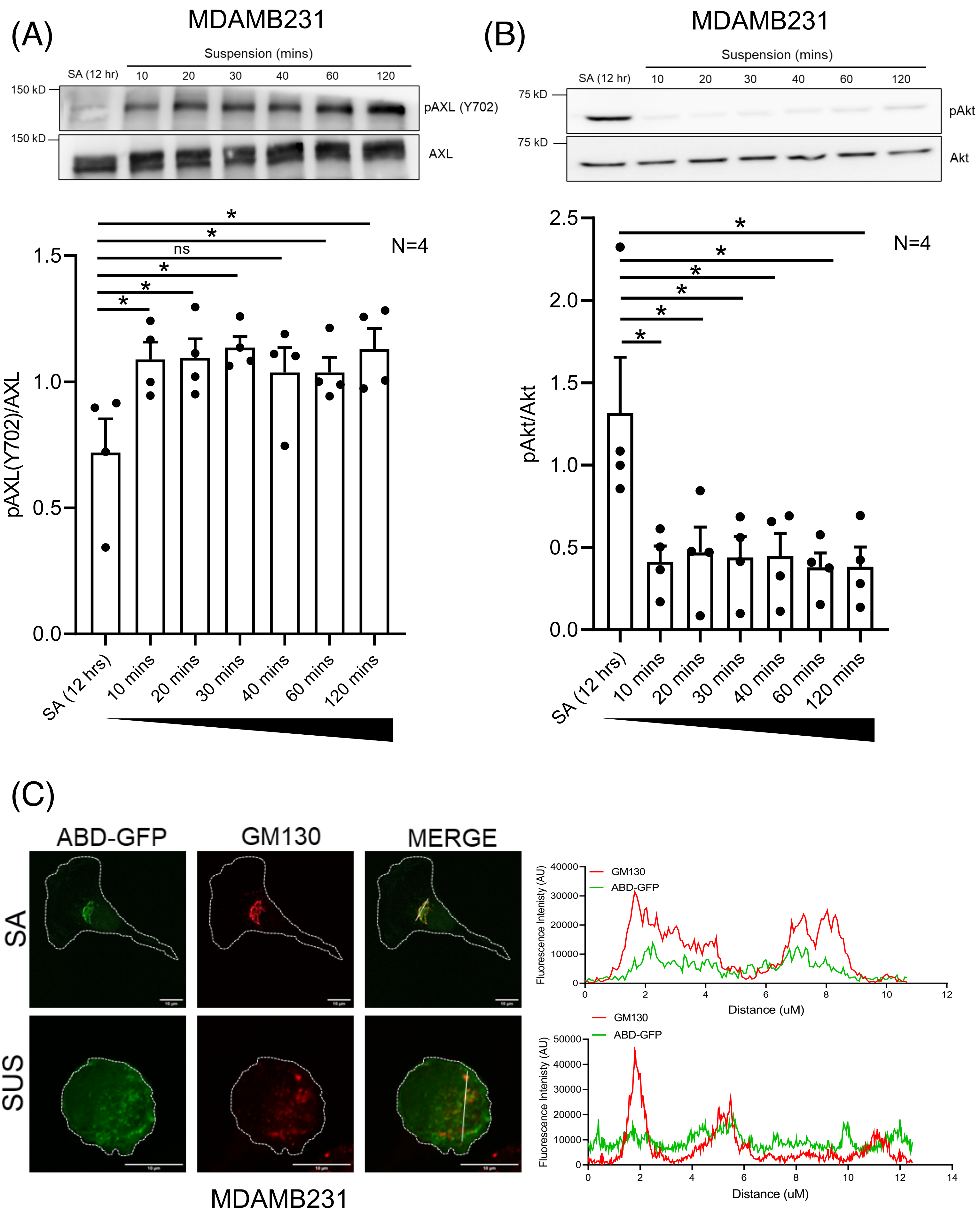

MDAMB231

(D)

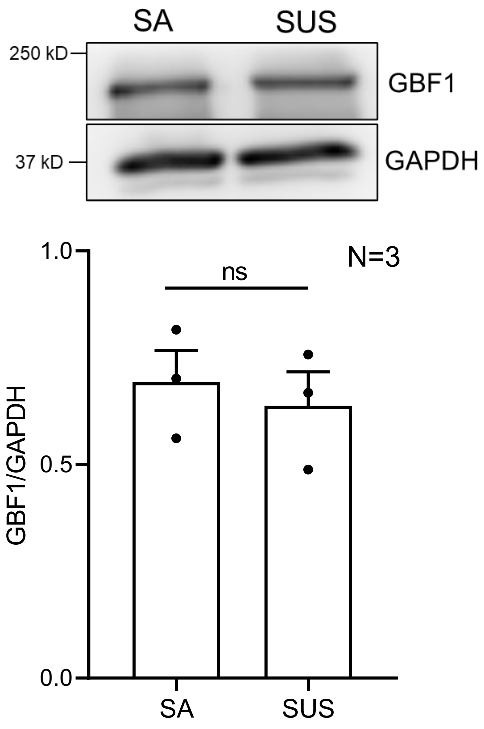

(A)

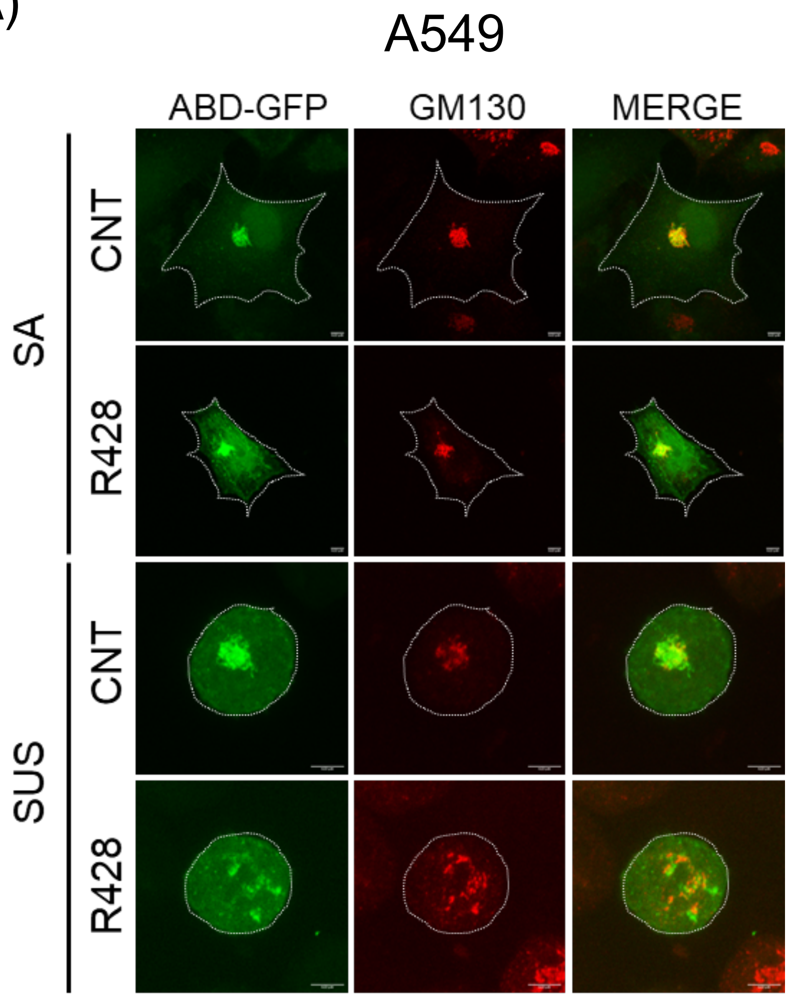
